# Different capsid-binding patterns of the β-herpesvirus-specific tegument protein pp150 (pM32/pUL32) in murine and human cytomegaloviruses

**DOI:** 10.1101/420604

**Authors:** Wei Liu, Xinghong Dai, Jonathan Jih, Karen Chan, Phong Trang, Xuekui Yu, Rilwan Balogun, Ye Mei, Fenyong Liu, Z. Hong Zhou

## Abstract

The phosphoprotein pp150 is a structurally, immunogenically, and regulatorily important capsid-associated tegument protein abundant in β-herpesviruses including cytomegaloviruses (CMV), but absent in α-herpesviruses and Γ-herpesviruses. In human CMV (HCMV), bridging across each triplex and three adjacent major capsid proteins (MCPs) is a group of three pp150 subunits in a “△”-shaped *fortifying* configuration, 320 of which encase and stabilize the genome-containing capsid. Because murine CMV (MCMV) has been used as a model for HCMV pathogenesis and therapeutic studies, one might expect that pp150 and the capsid in MCMV and HCMV have similar structures. Here, by cryoEM and sub-particle reconstructions, we have obtained structures of MCMV capsid and pp150 at near atomic resolutions and built their atomic models. Surprisingly, the capsid-binding patterns of pp150 differ between HCMV and MCMV despite their highly similar capsid structures. In MCMV, pp150 is absent on triplex Tc and exists as a “Λ”-shaped dimer on other triplexes, leading to only 260 groups of two pp150 subunits per capsid in contrast to 320 groups of three pp150 subunits encasing each HCMV capsid. Many more amino acids contribute to pp150-pp150 interactions in MCMV than in HCMV, making MCMV pp150 dimer inflexible thus incompatible to instigate triplex Tc-binding as observed in HCMV. While pp150 is essential in HCMV, pp150-deleted MCMV mutants remained viable though with attenuated infectivity and exhibiting defects in retaining viral genome. These results support targeting capsid proteins, but invalidate targeting pp150, when using MCMV as a model for HCMV pathogenesis and therapeutic studies.

**Importance:** CMV infection is a leading viral cause of congenital birth defects and often responsible for life-threating complications in immunocompromised individuals like AIDS and post-organ transplantation patients. Absence of effective vaccines and potent drugs against CMV infections has motivated animal-based studies, mostly based on the mouse model with MCMV, both for understanding pathogenesis of CMV infections and for developing therapeutic strategies. Here, we present the first atomic structures of MCMV and show that the organization patterns of capsid-associated tegument protein pp150 between human and mouse CMV are different despite their highly similar capsid structures. Our functional studies demonstrate that deleting pp150 does not eliminate MCMV infection in contrast to pp150’s essential role in HCMV infections. These results thus establish the validity to target capsid proteins, but raise concerns to target pp150, when using MCMV as HCMV model for pathogenesis and therapeutic studies.

## Introduction

Cytomegalovirus (CMV) is a member of the β-herpesvirus subfamily of *Herpesviridae* and can establish lifelong subclinical (latent) infection among the majority of the human population. Active human CMV (HCMV) infection is the leading viral cause of birth defects (*in utero* and in neonates) and often the culprit of life-threatening complications in immunocompromised individuals, such as organ transplant recipients and AIDS patients. Currently, there is no licensed vaccine against HCMV infection and conventional anti-HCMV drugs have well-known adverse side effects and are compromised by resistance (1). These factors call for novel approaches toward vaccine design and drug development against HCMV infections. Thanks to similarities in pathology and disease manifestation caused by CMV infections of humans and mice (*e.g.*, 2, 3-5, and review 6), pathogenesis studies and therapeutic developments have often relied on murine CMV (MCMV) as a model to evaluate the efficacy of lead compounds (7-9) and vaccine candidates (10, 11).

Recent high-resolution cryoEM structures of human herpesviruses (12-14), particularly the demonstration of inhibitors designed based on the structure of small capsid protein (SCP) (12, 13), have opened the door to structure-guided design of new drugs and vaccines targeting HCMV capsid proteins and the β-herpesvirus-specific tegument protein pUL32 (or phosphoprotein pp150, see review 15). The cryoEM reconstruction of HCMV at 3.9 Å resolution (14) reveals that pUL32 forms a unique capsid-binding tegument layer, likely to secure encapsidation of its dsDNA genome of 235 kbp, which is the largest among all herpesviruses. Particularly, 320 groups of three pUL32nt subunits form a “△”-shaped *fortifying* structure on every triplex. pUL32 is an abundant and immunogenic protein that is essential for HCMV virion egress and maturation (16, 17). Yet careful examination of their genomes suggests there might be structural differences between HCMV and MCMV. For example, HCMV pUL32 sequence is about 40% longer than pM32 (the homolog of pUL32 in MCMV) (18), suggesting that an examination of the structure of MCMV in detail may be fruitful in assessing the similarities and differences between HCMV and MCMV. Therefore, the functional and structural significance of pM32 remains to be established, in stark contrast to the large body of MCMV-based cell and animal studies concerning CMV infections.

Here, by cryoEM and sub-particle reconstructions, we have obtained structures of the MCMV capsid and its associated pM32 (pp150) at near atomic resolutions and built their atomic models, the first for any MCMV proteins. Comparison of the virion structures of MCMV and HCMV reveals that the patterns of pp150 binding to capsid differ between HCMV and MCMV, despite highly similar structures of their capsid proteins, including SCP. The atomic details underlying pp150-pp150 and pp150-capsid protein interactions rationalize the different capsid-binding patterns of pp150 in MCMV and HCMV. Our mutagenesis studies further establish pp150’s varying levels of functional significance in MCMV and HCMV infections. These results thus establish the validity of using MCMV as HCMV model for pathogenesis and therapeutic studies when targeting capsid proteins, but raise concerns when targeting pp150.

## Results

### Structures of MCMV capsid and capsid-associated tegument protein pM32

A technical challenge of determining the structure of herpesvirus particles is the enormous size of herpesvirus virions exceeding 200 nm in diameter, creating a focus gradient across the large sample thickness needed to fully embed the virion and the breakdown of the *Central Projection Theorem* due to a curved Ewald-sphere (14, 19). Initial cryoEM reconstruction from the 1,200 MCMV virion particle images (*e.g.*, Fig. S1A-D) recorded on a CCD camera was limited to ∼12 Å resolution (Fig. S1E-H). To alleviate the Ewald-sphere curvature effect (19, 20), we attempted to reduce the sample thickness by partially solubilizing the viral envelope through mild detergent treatment (Fig. S2A). From a total of 47,982 particles of MCMV virions from 2,200 300kV cryoEM micrographs recorded on photographic films (*e.g.*, Fig. S2A), we obtained an icosahedral reconstruction at ∼5 Å resolution (Fig. 1A, Movie S1). Local resolution assessment by *Resmap* (Fig. S2B) (21) indicates that densities of the capsid shell and especially near the base of the capsomers have the best resolution (from 4.0 Å to 4.5 Å) and that densities at the outmost radii and inside the capsid shell (*i.e.*, DNA genome) have resolutions worse than 4.5 Å, likely due to a combination of structural heterogeneity/flexibility (for DNA-related densities) and the more severe Ewald-sphere curvature effect for densities at larger radii.

**FIG. 1.**
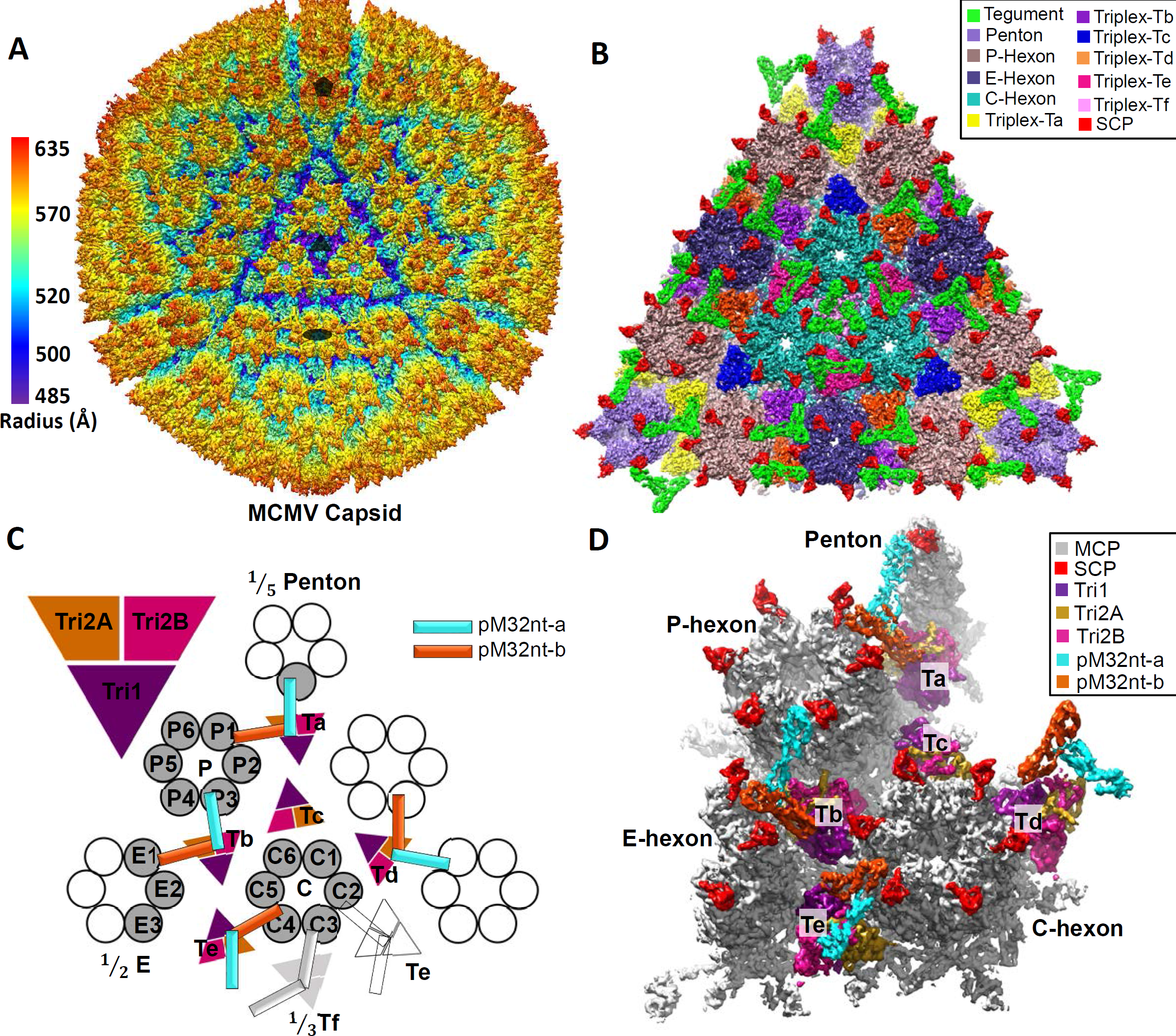
The 3D reconstruction of MCMV icosahedral capsid. (**A**) Radially colored surface representation of the 3D reconstruction of MCMV at ∼5 Å resolution, viewed along a 3-fold axis. Only the icosahedral symmetric components, including the capsid and capsid-associated tegument complex (CATC), are visible because the pleomorphic envelope and tegument layer, as well as the unique portal complex, have been smeared due to averaging of tens of thousands of particles and imposition of icosahedral symmetry. The filled pentagon, triangle and oval mark a 5-fold, 3-fold, and 2-fold axis, respectively. (**B**) Zoom-in surface view of one facet of the icosahedral capsid with structural components colored as indicated. Density of triplex Tf region at the center is smeared due to imposition of 3-fold symmetry during icosahedral reconstruction. A slightly lower density threshold was used for the pentons and triplex Ta/Tf with their associated tegument protein pM32, such that their volumes are comparable to subunits elsewhere. (**C-D**) Schematic (C) and cryoEM density map (D) of one asymmetric unit (shaded) of the reconstruction with individual capsid protein subunits labeled, following the nomenclature used in HCMV (14). Subunits of triplex Tf and its associated pM32 are shown in light gray instead of colors as their orientations were not determined in the icosahedral reconstruction here, but from sub-particle reconstruction after sub-particle 3D classification (see Fig. S5).

Though at different resolutions, both the early reconstruction from CCD images of intact virions and the higher resolution reconstruction from detergent-treated virions show identical structural organizations of tegument and capsid proteins (Fig. S1E and Fig. 1A). The ∼5 Å 3D reconstruction shows a highly conserved T=16 icosahedral virion capsid revealing the molecular boundaries among 12 pentons, 150 hexons, 320 triangular triplexes and 260 pM32 (pp150) dimers (Fig. 1A-B), allowing identification of individual molecules (Fig. 1C-D). Imposing icosahedral symmetry during 3D reconstruction both weakens the density and lowers the resolution of regions with deviation from strict icosahedral symmetry. One of the twelve icosahedral vertices of the herpesvirus capsid does not contain a penton, but a DNA packaging/ejection portal complex (22-25); thus, tegument densities interacting with pentons are weaker than those interacting only with hexons. The 3-fold symmetrized tegument densities associating with triplex Tf are also weakened and are only visible when displayed at a lower density threshold (see Fig. S5 for a sub-particle reconstruction of this region showing a pp150 dimer attached to triplex Tf).

The structural components within an asymmetric unit encompass 1/5 of a penton capsomer, 2.5 hexon capsomers (1 P hexon, 1 C hexon, and 1/2 of an E hexon), 5 and 1/3 triplexes/pM32 dimers (Ta, Tb, Tc, Td, Te, and 1/3 Tf) (Fig. 1C). To overcome the aforementioned problem of weakened densities and limited resolutions, we have used a sub-particle reconstruction strategy to obtain structures of the MCMV capsid and its associated pM32 at near atomic resolutions (Fig. 2B-E, Fig. S3-S5, Movie S2-S4) and also built their atomic models (Fig. S6-S9). In particular, our sub-particle reconstruction of the region around the 3-fold axis shows that pM32 on triplex Tf also exists as a dimer (Fig. S5), as with triplexes Ta, Tb, Td, and Te (Fig. 1A-D). The improved resolution of the sub-particle reconstructions allowed us to build *de novo* atomic models for the N-terminal one-third portion of pM32 (pM32nt) and the four capsid proteins [the major capsid protein (MCP), the small capsid protein (SCP), the triplex monomer protein (Tri1), the triplex dimer protein (Tri2)]. In total, we have built atomic models of 55 unique conformers, including 16 MCP, 16 SCP, 5 Tri1, 5 Tri2A, 5 Tri2B and 8 pM32nt (Fig. 2H), amounting to over 27,000 amino acid residues. The atomic models of the MCMV capsid proteins are highly similar to those in HCMV (14), but those for pM32 and pUL32 differ, as detailed below.

**FIG. 2.**
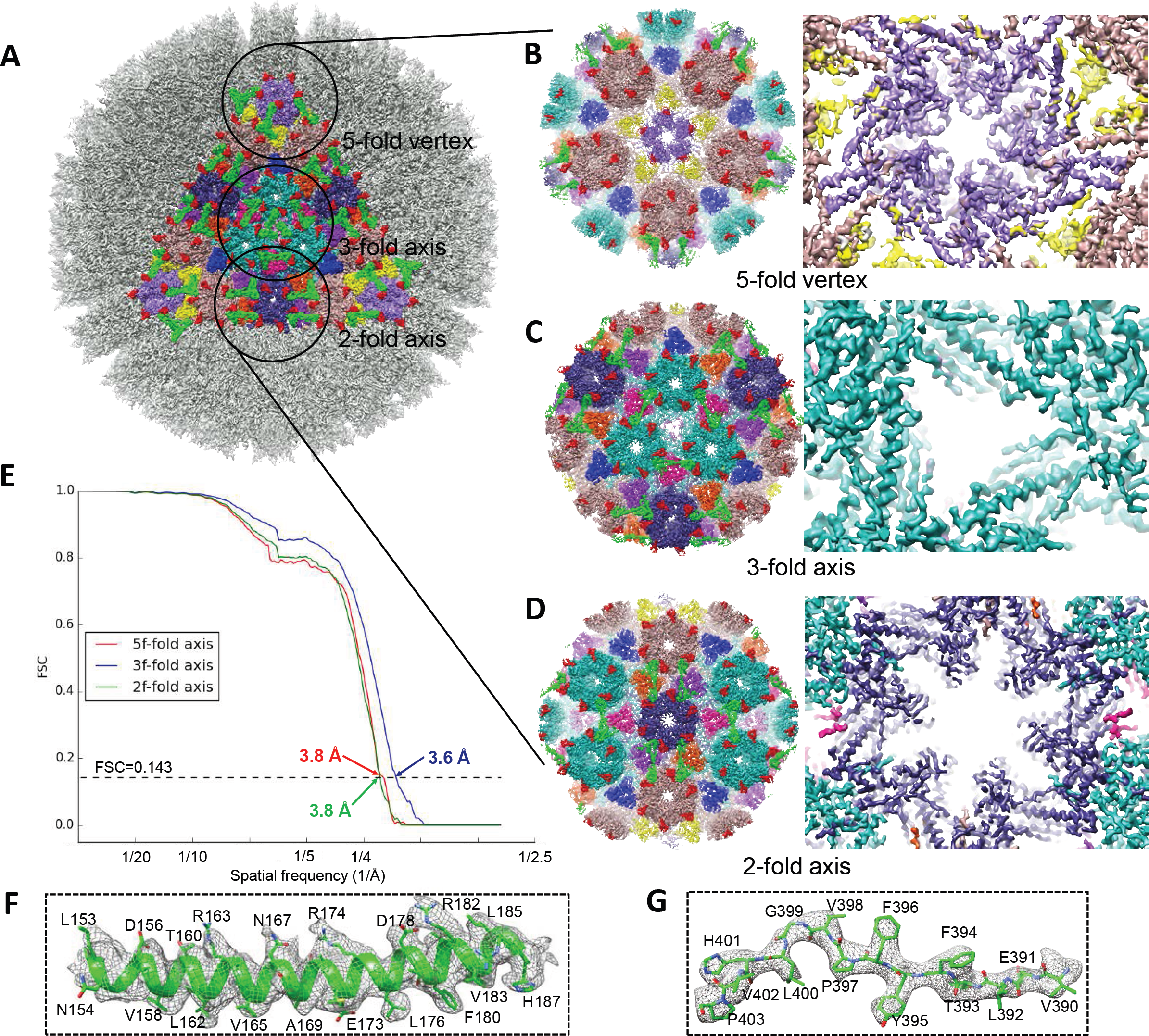
Resolution assessment, sub-particle reconstructions and MCMV atomic models. (**A-D**) The icosahedral reconstruction in Figure 1A is reshown (A) with one triangular facet in color and regions (referred to as “sub-particles”) surrounding a 5-fold, 3-fold, and 2-fold axis circled. A total of 575,784 5-fold sub-particles, 959,640 3-fold sub-particles, and 1,439,460 2-fold sub-particles were boxed out from original particle images and refined to yield improved resolutions for the 5-fold sub-particle (B), 3-fold sub-particle (C), and 2-fold sub-particle (D). The enlarged views from inside (right panels) of these sub-particle reconstructions show α-helices and β-strands with well-resolved side chain densities. (**E**) Gold-standard (0.143) Fourier shell correlation (FSC) curves of the sub-particle reconstructions indicating the resolutions of sub-particle reconstructions at the 5-fold (red), 3-fold (blue), and 2-fold (green) axes are 3.8 Å, 3.6 Å, and 3.8 Å, respectively. (**F-G**) Close-up views of cryoEM density map (gray mesh) of an α-helix (F) and a loop (G) in the MCP floor region, superposed with atomic models (color). (**H**) *De novo* atomic models of individual capsid (MCP, SCP, Tri1, Tri2A, and Tri2B) and tegument (pM32nt) proteins shown as rainbow-colored ribbons (blue at the N-terminus to red at the C-terminus).

### Atomic models of MCP and SCP, and their interactions in penton and hexons

An MCP monomer binds an SCP monomer to form a heterodimer (Fig. 4B), which constitutes one of the five and six subunits in penton and hexon capsomers, respectively (Fig. 4C). Like that of HCMV (14), the structure of the 1,353 a.a. long (149 kDa) MCP monomer of MCMV consists of seven domains (Fig. 3A): upper (a.a. 477-1015), channel (a.a. 398-476, 1303-1353), buttress (a.a. 1090-1302), helix-hairpin (a.a. 190-233), dimerization (a.a. 291-362), N-lasso (a.a. 1-59), and a bacteriophage HK97-like (“Johnson”)-fold (a.a. 60-189, 234-290, 363-397, 1016-1089). These domains are located in the tower (upper, channel, and buttress domains) and the floor (helix-hairpin, dimerization, N-lasso, and Johnson-fold domains) regions of each capsomer subunit (Fig. 2H). Because the Johnson-fold domain of MCMV MCP corresponds to the entire molecule of the HK97 MCP (gp5) (26), segments corresponding to HK97 gp5’s four domains—axial (A), extended loop (E loop), peripheral (P), and spine helix—are designated as A, E-loop, P, and spine helix sub-domains, respectively (Fig. 3D).

**FIG. 3.**
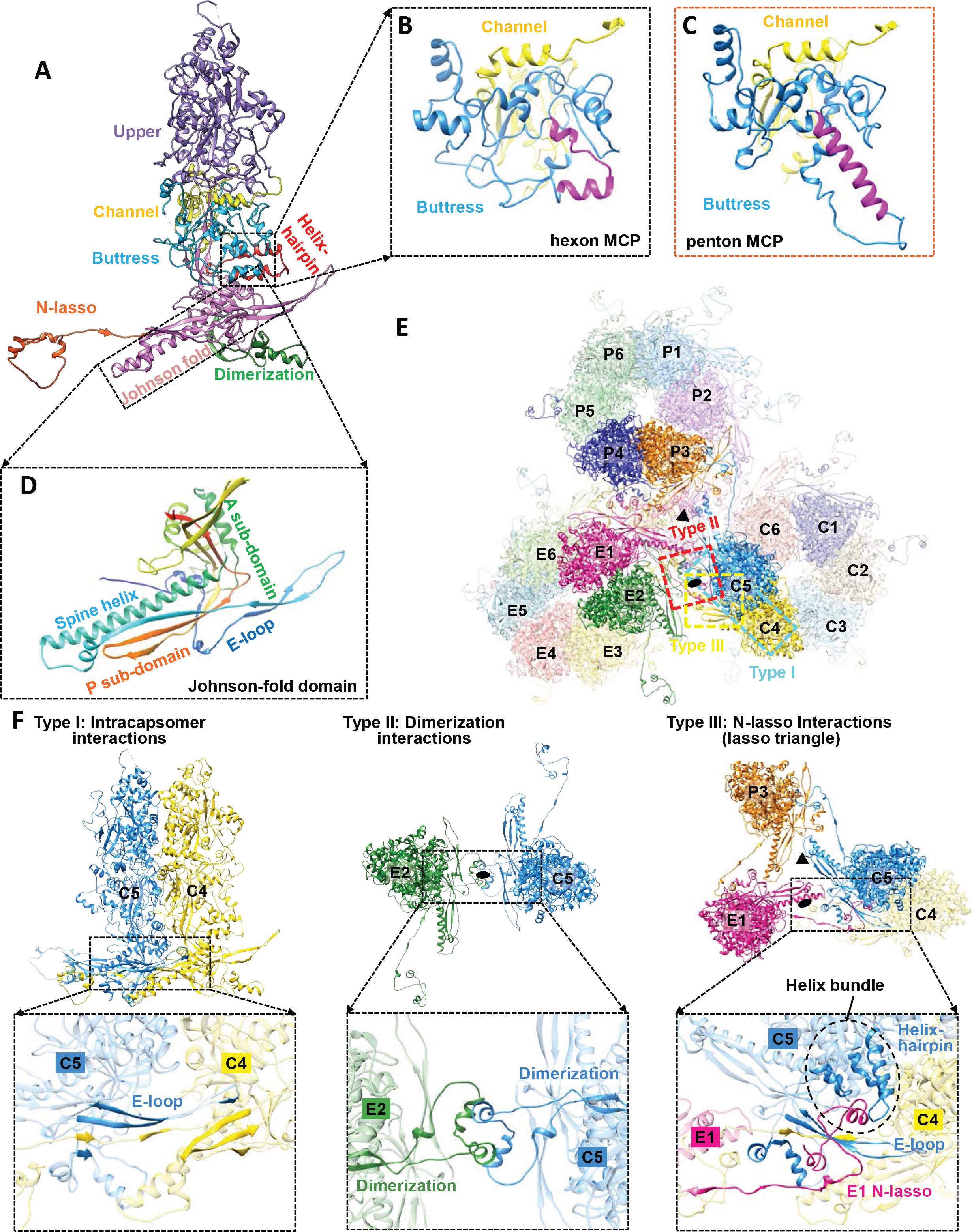
MCP structure and three types of capsid floor-defining MCP-MCP interactions. (**A**) Domain organization in the hexon MCP. (**B-C**) Comparison of the buttress domain in hexon (B) and penton (C) MCP. An elbow-like helix-turn-helix structure (magenta in B) in the buttress domain of hexon MCP is folded into a single long helix (magenta in C) in penton MCP. (**D**) The structure of the Johnson-fold domain shown in rainbow-colored ribbon with sub-domains labeled. (**E**) Overview of MCPs from C, E, and P hexons. Local 3-fold and 2-fold axes are indicated by a triangle and an oval, respectively. Three major types of MCP-MCP interactions are boxed in cyan (type I), red (type II), and yellow (type III), respectively. (**F**) Details of the three types of MCP-MCP interactions.

**FIG. 4.**
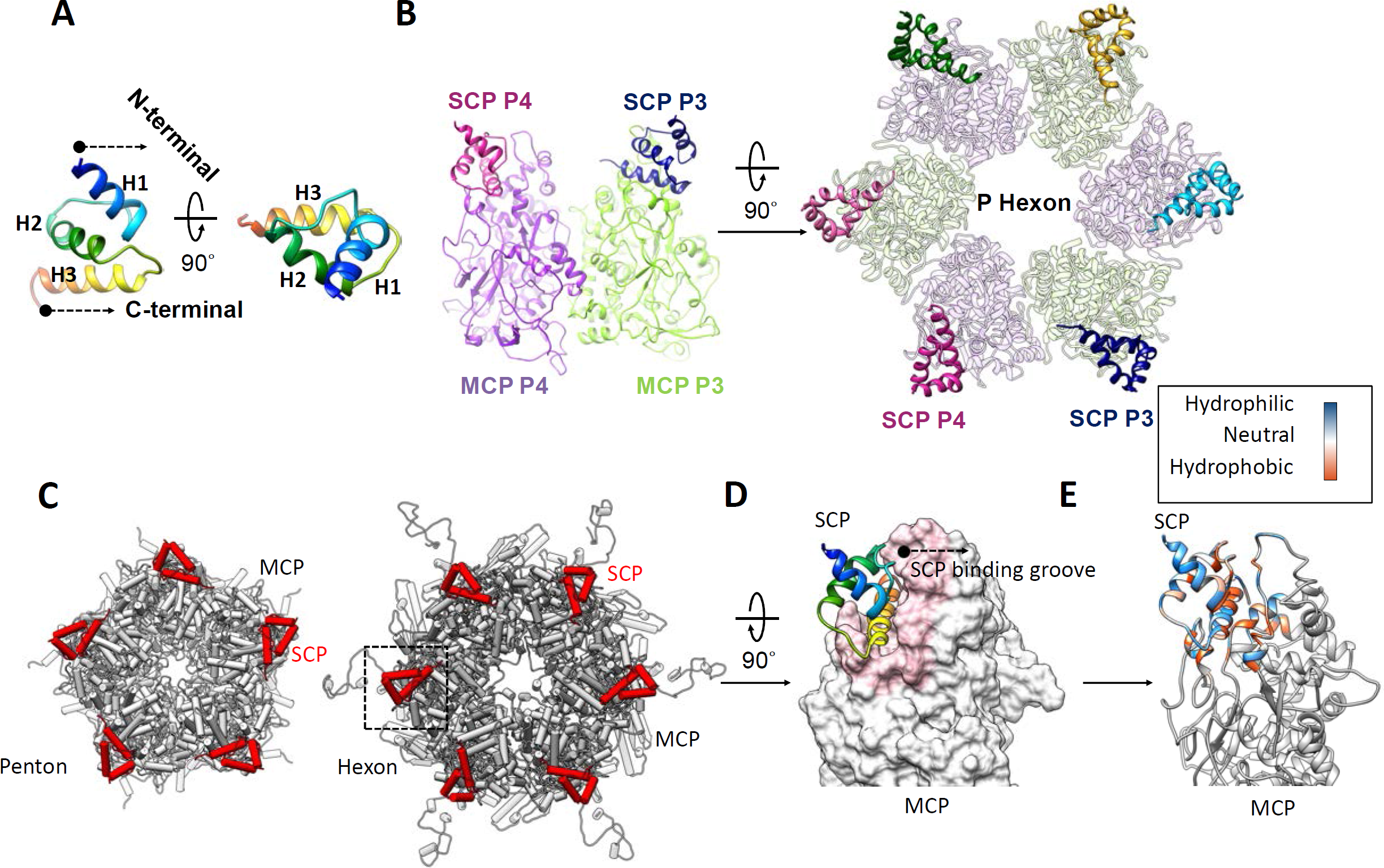
MCP interactions with SCP. (**A**) SCP shown as rainbow-colored ribbon in two orthogonal reviews. (**B**) SCP binding to the upper domain of hexon MCP, as seen from the side of two adjacent MCP subunits (left panel) and from the top of a P hexon (right panel). Only MCP upper domains are shown. (**C**) Pipe-and-plank depictions of a penton (left panel) and hexon (right panel) with SCP and MCP subunits shown in red and gray, respectively. (**D-E**) SCP (ribbons) bound to the MCP upper domain, shown either as surface (D) with the SCP-binding groove highlighted in pink, or as ribbons with the hydrophobicity properties of the structures around the SCP-binding groove in color (E). SCP-MCP interactions are mainly hydrophobic.

MCP monomers in hexons and pentons have distinct conformations. For instance, the sequence segment corresponding to the helix-loop-helix motif in the buttress domain of hexon MCP (Fig. 3B) folds into a single long helix in penton MCP (Fig. 3C). As in HCMV (14), there are three notable types of network interactions in the MCP floor regions (Fig. 3E-F). Type I interactions are intra-capsomeric β-sheet augmentations which occur between two adjacent MCPs within a capsomer. As exemplified by C4 and C5 MCPs in Figure 3F (left panel), two β-strands from the E-loop of Johnson-fold domain and one β-strand from the dimerization domain of C5 MCP are joined by two β-strands from the N-lasso domain of C4 MCP, resulting in a five-stranded β-sheet (Fig. 3F, left panel). Type II interactions are inter-capsomeric interactions that occur between two pairs of α-helices in the dimerization domains of MCPs across local 2-fold axes, as illustrated by E2 and C5 MCPs in Figure 3F (middle panel). Like type II interactions, type III interactions are inter-capsomeric interactions, but are formed by three MCPs featuring the lassoing action of the N-lasso domain (E1), which extends out and lashes around an E-loop (C5) and a N-lasso neck (C4) of two type I interacting MCPs located across a local 2-fold axis (Fig. 3F, right panel). Additionally, a small helix bundle is formed from an α-helix from E1 N-lasso, two α-helices from the helix-hairpin domain of C5 MCP, and an α-helix from the buttress domain of C5 MCP, further securing the E1 N-lasso (Fig. 3F right panel).

Our model of SCP encompasses residues 36-95 of the 98 a.a. long M48.2 gene product and consists of three 3.5-turn α-helices and two connecting loops, folded into a triangular spiral with the N-terminal helix (H1) pointing outwards (Fig. 4A). The H3 helix of SCP inserts into a deep groove in a region of MCP upper domain rich in α-helices and loops (Fig. 4D). Sequence-based surface analysis indicates that predominantly hydrophobic interactions contribute to MCP-SCP binding (Fig. 4E).

### Atomic models of the triplexes

Each triplex is a heterotrimer containing one unique conformer of Tri1 and two conformers of Tri2—Tri2A and Tri2B—that “embrace” each other to form a dimer (Fig. 5G). Tri2 monomer consists of three domains: clamp (a.a. 1-88), trunk (a.a. 89-183, 291-311), and embracing arm (a.a. 184-290). While the clamp and trunk domains in Tri2A and Tri2B are nearly identical (RMSD is only 1.12 Å) and are superimposable by a ∼120^°^ rotation about the local 3-fold axis (Fig. 5F, right panel), their embracing arms differ by a ∼45^°^ bend with an RMSD of 3.69 Å (Fig. 5H).

**FIG. 5.**
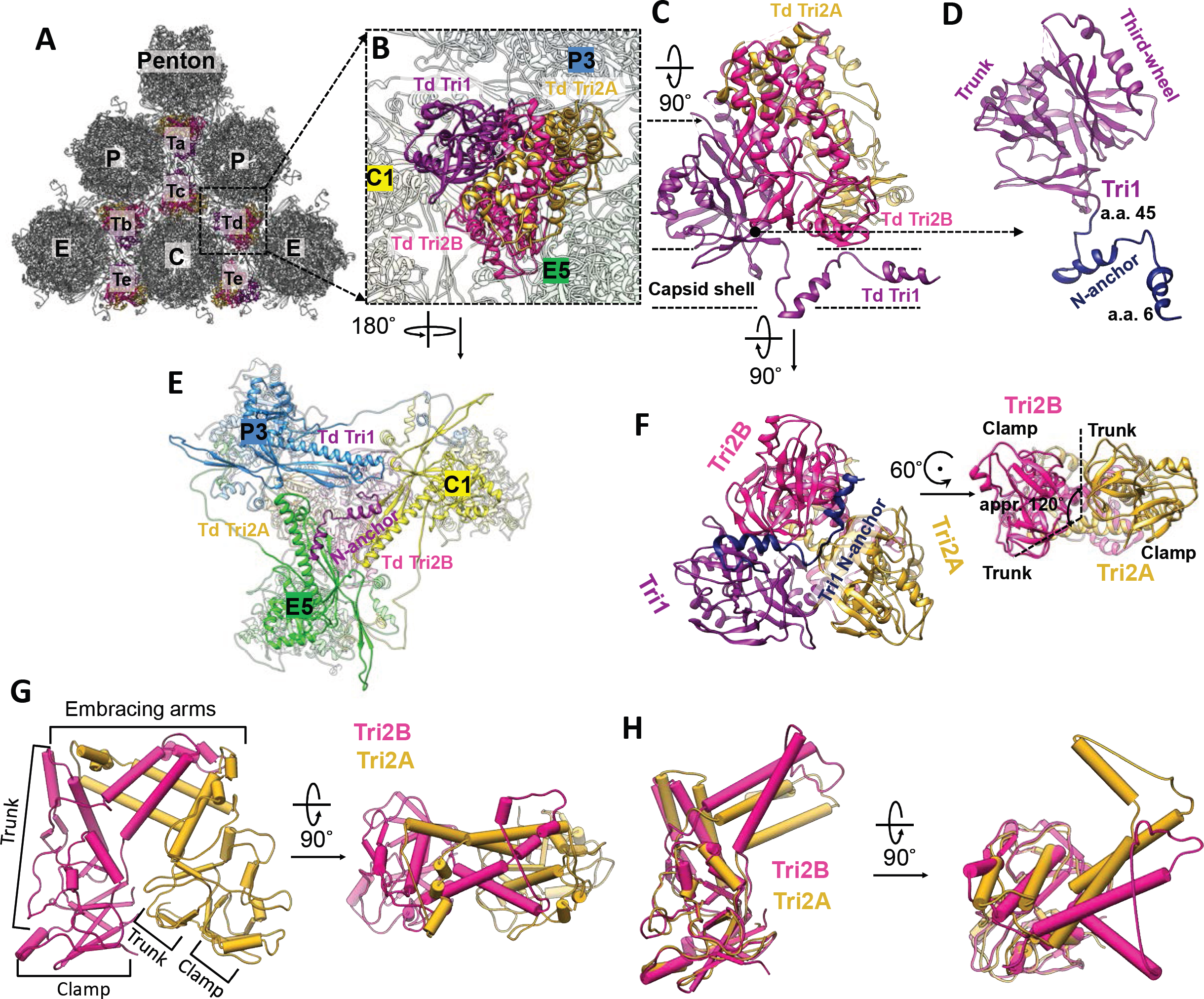
Structure of triplex and functional significance of Tri1 N-anchor. (**A**) Distribution of triplexes Ta, Tb, Tc, Td, and Te among penton and three types of hexons (C, E, and P). (**B**) Enlarged view of a triplex Td with three adjacent hexon MCP subunits labeled (C1, E5, and P3). (**C-D**) Details of the structures of triplex Td (C) and Tri1 (D). (**E**) Bottom view of (B) showing that triplex Td anchors to the capsid floor by the “V”-shaped Tri1 N-anchor. For clarity, only part of MCP subunits C1, E5, and P3 is shown. (**F**) Triplex Td viewed from inside of the capsid showing similar clamp and trunk domains, albeit rotated about 120^°^ relative to each other. (**G**) Pipe-and-plank representations of Tri2 dimer in side (left panel) and top (right panel) views showing the helix bundle formed from Tri2A and Tri2B’s embracing arm domains. (**H**) Superposition of Tri2A and Tri2B showing nearly identical clamp and trunk domains, but different embracing arms.

Clung to the side of the two *embracing* Tri2 subunits is the Tri1 monomer, which consists of three domains: N-anchor (a.a. 1-45), trunk (a.a. 46-171), and third-wheel (a.a. 172-294) (Fig. 2H, Fig. 5D). The helix-loop-helix-loop motif of the N-anchor domain traverses across the capsid floor near its local 3-fold axis such that both its helices fill the valley between the P-subdomain β-sheet and the spine helix of a Johnson-fold domain of an MCP subunit (Fig. 5C and 5E, Movie S1). Thus, N-anchor *anchors* Tri1 and the entire triplex from inside the capsid beneath the MCP floor, simultaneously sealing the hole at the local 3-fold axis where three neighboring MCP P-subdomains assemble. Notably, such “internally anchored” interactions could be pressure-fortified from within the capsid by DNA during genome packaging (12).

### MCMV and HCMV have distinctive capsid-binding patterns of pM32 and pUL32

The reconstruction densities exhibit different patterns of tegument-capsomer association between MCMV and HCMV (Fig. 6A and 6B). Our MCMV reconstructions show 260 tegument densities (Fig. 6A), as opposed to the 320 tegument densities of HCMV (Fig. 6B) (14, 27). In addition, their detailed structures differ, existing as a group of two subunits in MCMV (Fig. 6A) and a group of three in HCMV (Fig. 6B), with arrangements resembling the Greek alphabets “Λ” and “△”, respectively. Moreover, as shown in yellow in Figure 6A, pM32 does not bind to triplex Tc in the MCMV capsid. Neither are pM32 density connections observed between edge and facet capsomers. Specifically, pM32 dimers of the edge type join P hexons, E hexons, (Fig. S11) and pentons on the 20 edges of the icosahedral capsid to triplexes Ta, Tb, and Td (Fig. 1B, Fig. 6A), while pM32 dimers of the facet type bind three C hexons together in the center of each icosahedral facet to triplexes Te and Tf (Fig. 1B, Fig. 6A, Fig. S11). In contrast, all triplexes of the HCMV nucleocapsid—including triplexes Tc—are occupied by three pUL32 subunits arranged as a dimer and a monomer of pUL32. Atop each triplex, dimeric pUL32 branches out to interact intimately with their closest respective capsomer subunits, in analogous fashion to the “Λ”-shaped tegument densities in MCMV (Fig. 6B). However, the third (monomeric) copy of pUL32 in HCMV bridges the top of the triplex with a third neighboring capsomer (Fig. 6B). This monomeric tegument density, in conjunction with the presence of pUL32 subunits above triplex Tc, facilitates connections between the edge and facet capsomers of HCMV (Fig. 6B) (27) *not* observed analogously in MCMV.

**FIG. 6.**
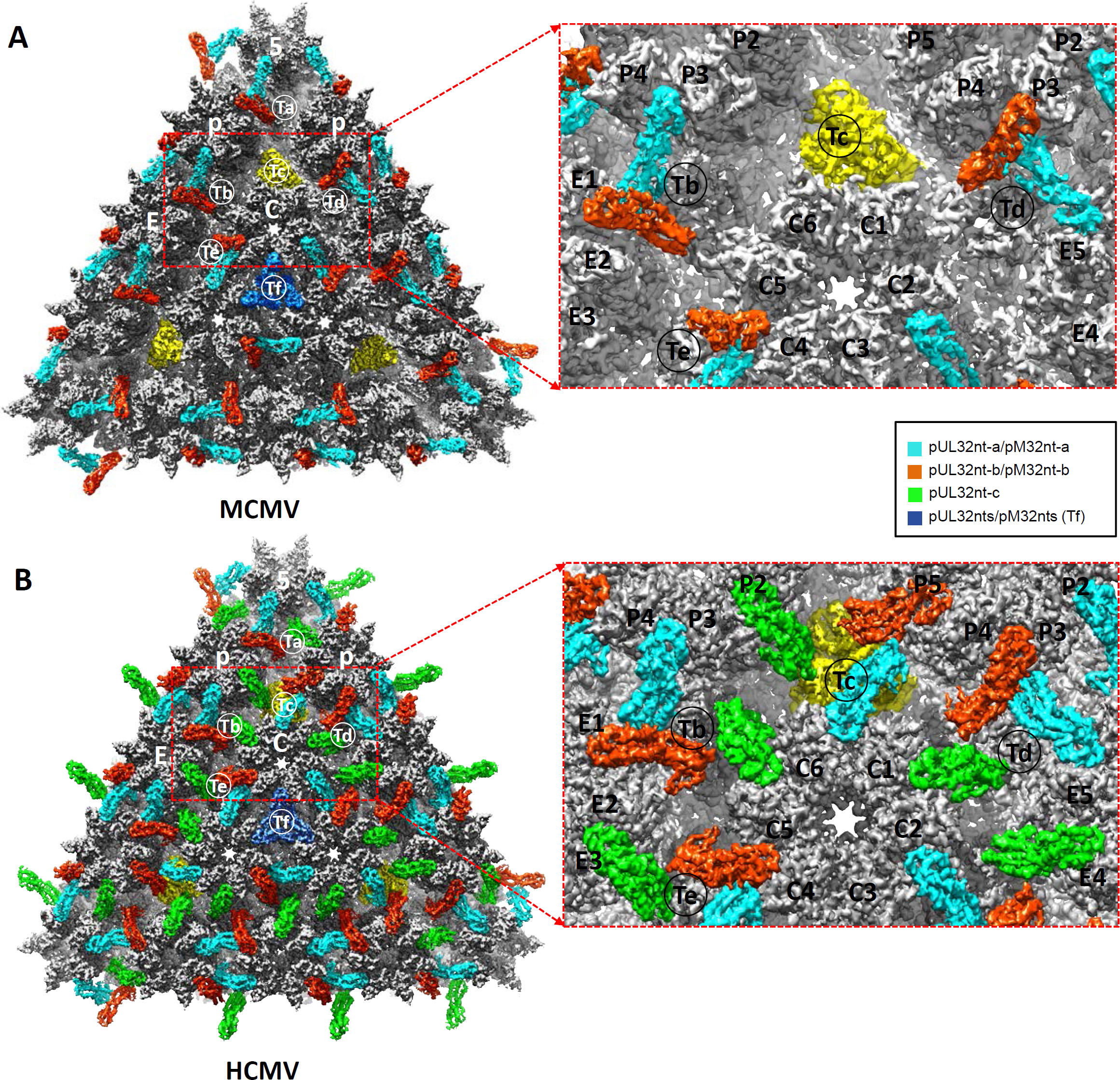
Comparison of capsid-binding patterns of CATC in MCMV and HCMV. (**A**) A triangular facet of MCMV icosahedral reconstruction. Except for triplex Tc (highlighted in yellow), two pM32nt subunits form a “Λ”-shaped density (cyan and orange red) acts like a stayed cable, with each of its two arms holding to a neighboring hexon/penton capsomer and its vertex sitting atop a triplex. (**B**) A triangular facet of HCMV icosahedral reconstruction (14). Three pUL32nt subunits form a “△”-shaped *fortifying* structure on every triplex, two of which (cyan and orange red) form a “Λ”-shaped structure similar to that in MCMV, and the third (green) bridges the gap between edge and facet capsomers. The density of pM32nt/pUL32nt in Tf region was colored in blue as the densities in this region were smeared after imposing 3-fold symmetry during icosahedral reconstruction (See how this has been resolved in sub-particle reconstruction in Fig. S5).

### Structure of pM32 and its dimer interface

Our atomic models of MCMV pM32 contain an N-terminal of approximately one-third of full-length (718 a.a.) pM32 (pM32nt-a, consisting of a.a. 66-87, 91-105, 137-150, 173-295; pM32nt-b, consisting of a.a. 66-113, 128-156, 173-295). Similar to pUL32nt of HCMV, pM32nt of MCMV is dominated by α-helices (Fig. 7A, Fig. S15) and characterized by upper and lower helix bundles joined by a central long helix (∼69 Å in length, a.a. 208-253) (Fig. 2H, Fig. 7A). We also identified the conserved region 1 (CR1) and region 2 (CR2) in MCMV pM32nt (Fig. 7A). However, only one cysteine, rather than four, was identified in pM32’s equivalent sequence of HCMV pUL32’s cys tetrad (Fig. 7A) (18). All bound pM32s are dimerized in MCMV and can be classified into two types, either pM32-a (cyan) or pM32-b (red orange), based on their relative locations on the capsid (Fig. 1C-D). The subunits of pM32nt dimers cluster on each triplex and lean against two neighboring MCPs (Fig. S10A). pM32nt-a and pM32nt-b are similar in structure (the magnitude of RMSD is 1.20 Å) (Fig. 7B) and form a “Λ”-shape configuration. As shown in Figure S10B-C, the two conformers interact with capsid proteins via hydrophobic and/or hydrophilic interactions, bearing a marked resemblance to those in HCMV (Fig. S10E-F).

**FIG. 7.**
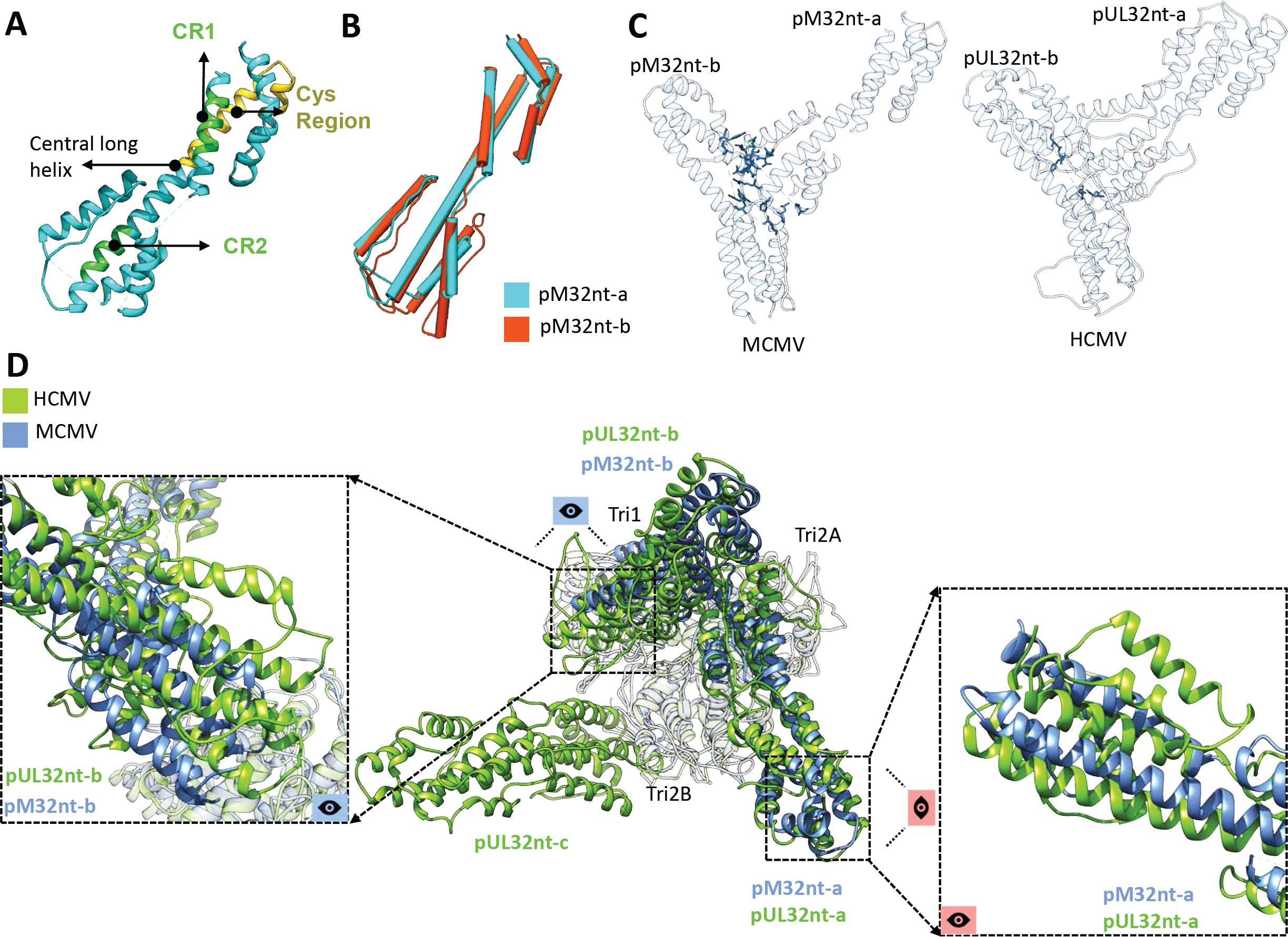
CATCs in MCMV and HCMV have conserved domain structures but divergent interacting interfaces. (**A**) Green residues denote β-herpesvirus-conserved regions CR1 and CR2, while yellow residues denote the primate CMV-conserved cys region in pM32nt. (**B**) Structural alignment based on the Cα atoms of the two pM32nt conformers reveals a large degree of structural similarity between pM32nt-a and pM32nt-b in MCMV. (**C**) Residues involved in MCMV pM32nt-pM32nt (left panel) and HCMV pUL32nt-pUL32nt (right panel) interactions (whose atoms are within 3 Å from each other, as exemplified by pp150 dimers from triplex Te regions in MCMV and HCMV) are highlighted in blue, respectively. (**D**) The atomic models of triplex Td (gray) in HCMV and MCMV were aligned as a rigid body to show how their associated CATCs differ. Conformers b of CATC (*i.e.*, pM32nt-b of MCMV and pUL32nt-b in HCMV) show major translational and rotational displacements (left panel) while conformers a (pM32nt-a and pUL32nt-a) only show minor rotational displacements (right panel) between HCMV and MCMV. Conformer c only exists in HCMV.

Careful comparison of the atomic model of pM32 dimer and the corresponding pUL32 subunits in HCMV (for example, pM32 and pUL32 dimer from triplex Te regions) reveals that 17 residues in the pM32-pM32 interface are within 3 Å of each other, as opposed to only 4 residues at the pUL32-pUL32 interface (Fig. 7C). The existence of 13 additional residues at the molecular interface of the pM32 dimer indicates a stronger and more rigid pM32-pM32 association than between pUL32-pUL32, which is congruent with the pairwise presence/absence of pM32 subunits on MCMV capsid. This also supports the observation that pM32 exists only as a dimer in our reconstructions in contrast to the existence of both monomeric and dimeric forms of pUL32 in HCMV (14).

Additionally, rigid-body fitting of the HCMV triplex-pUL32nt atomic model (for example, Td region) into MCMV triplex density (in order to align triplex models from MCMV and HCMV) reveals further comparative insights into pM32/pUL32 binding in MCMV and HCMV (Fig. 7D). First, as mentioned above, only two pM32 subunits bind with MCMV triplex as a dimer instead of three pUL32 subunits for each HCMV triplex. Second, the organization of pM32/pUL32 dimers in MCMV and HCMV with respect to triplex resemble each other, though the specific orientation of dimer subunits possesses some distinctions (Fig. 7D). In contrast to the minor rotational displacements exhibited between pM32-a and pUL32-a, pM32-b and pUL32-b show greater translational and rotational displacements (Fig. 7D). Third, Tri1 residues of MCMV and HCMV that interact with pM32 and pUL32, respectively, are conserved (Fig. S12) and conceivably play a crucial role in pM32/pUL32 binding.

In contrast to the binding of a pM32 dimer to each of triplexes Ta, Tb, Td, Te, and Tf, no pM32 is bound to triplex Tc (Fig. 6A, Fig. 8A). Rigid-body fitting of triplex Td decorated with pM32nt dimer into triplex Tc density (Fig. 8B) reveals that distances between pM32’s upper domains to their adjacent SCPs are greater at triplex Tc than triplex Td (7 Å vs. 4 Å for pM32nt-a; 14 Å vs. 6 Å for pM32nt-b) (Fig. 8C, left panels). The rigidity of pM32 dimer discussed above conceivably prevents pM32 from extending, or “spreading,” to span such long distances, explaining the absence of pM32 above triplex Tc in MCMV.

**FIG. 8.**
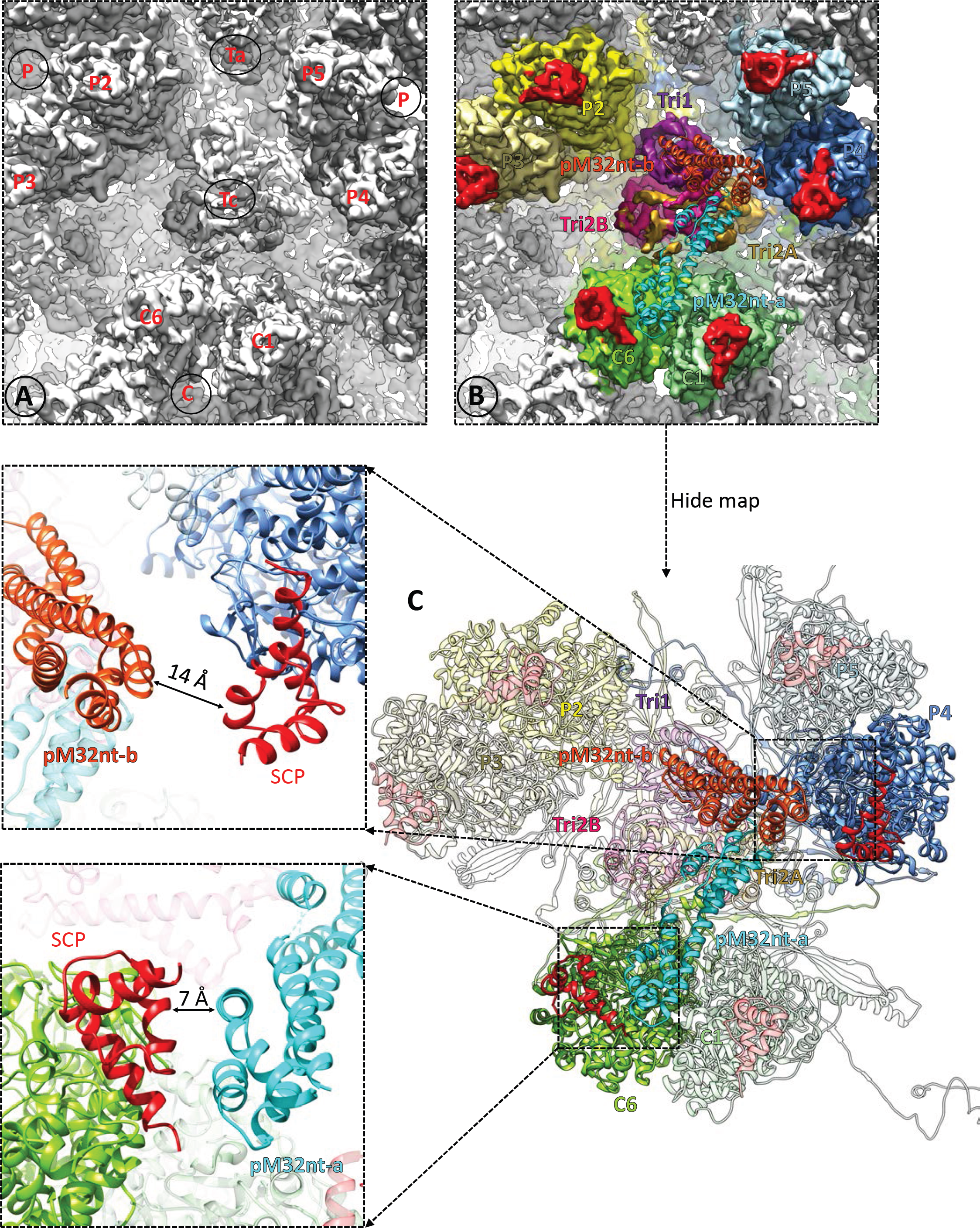
pM32 does not bind to triplex Tc in MCMV. (**A**) Zoom-in view of MCMV triplex Tc region. (**B**) The same region as shown in (A). Models of triplex and two pM32nt subunits from MCMV triplex Td region were docked as a rigid body into MCMV Tc region density by only fitting triplex structures into the map. Neighboring MCPs are colored individually while Tri1, Tri2A, Tri2B, pM32nt-a, pM32nt-b, and SCP are colored as in Figure 1D. (**C**) The right panel shows the atomic models which are presented using the same color scheme as in (B). The closest distances between pM32nt-a/pM32nt-b and their associated SCPs are shown in the left panels.

### MCMV mutant with deletion of the M32 sequence is attenuated in growth but is viable

Next, we show that pM32 is important, though not essential, for MCMV replication *in vitro* by successfully generating an infectious MCMV mutant with deletion of the coding sequence of M32, which we term ΔM32. To generate ΔM32, we adopted our previously-published protocols for generating gene-deletion mutants of HCMV to mutagenize a BAC clone of the wild-type MCMV (Smith strain) genome (MCMV_BAC_) by deleting the M32 open reading frame (28-30). Furthermore, rescued viral mutant, R-M32, was generated from ΔM32, by restoring the M32 sequence, following the procedures as described previously (29, 31). Deletion of the M32 gene in ΔM32 and the restoration of the M32 sequence in R-M32 were confirmed by PCR and Southern blot analysis.

To determine whether ΔM32 has any growth defects *in vitro* and whether pM32 is essential for MCMV replication in cultured cells, we measured the growth rates of mutant ΔM32, rescued mutant R-M32, and parental MCMVBAC viruses in NIH 3T3 cells. ΔM32 exhibited growth defective phenotype as the titers of ΔM32 were lower than those of MCMV_BAC_ in a 6-day growth study (Fig. 9A). At day 4, the titer of ΔM32 was about 100-fold lower than that of MCMV_BAC_ (Fig. 9A). The observations that ΔM32 grew in NIH3T3 cells indicate that M32 is not required for MCMV replication *in vitro*. Thus, in contrast to its HCMV homolog pUL32, which is essential for HCMV replication (29, 31), M32 is not essential—though important—for MCMV replication *in vitro*.

**FIG. 9.**
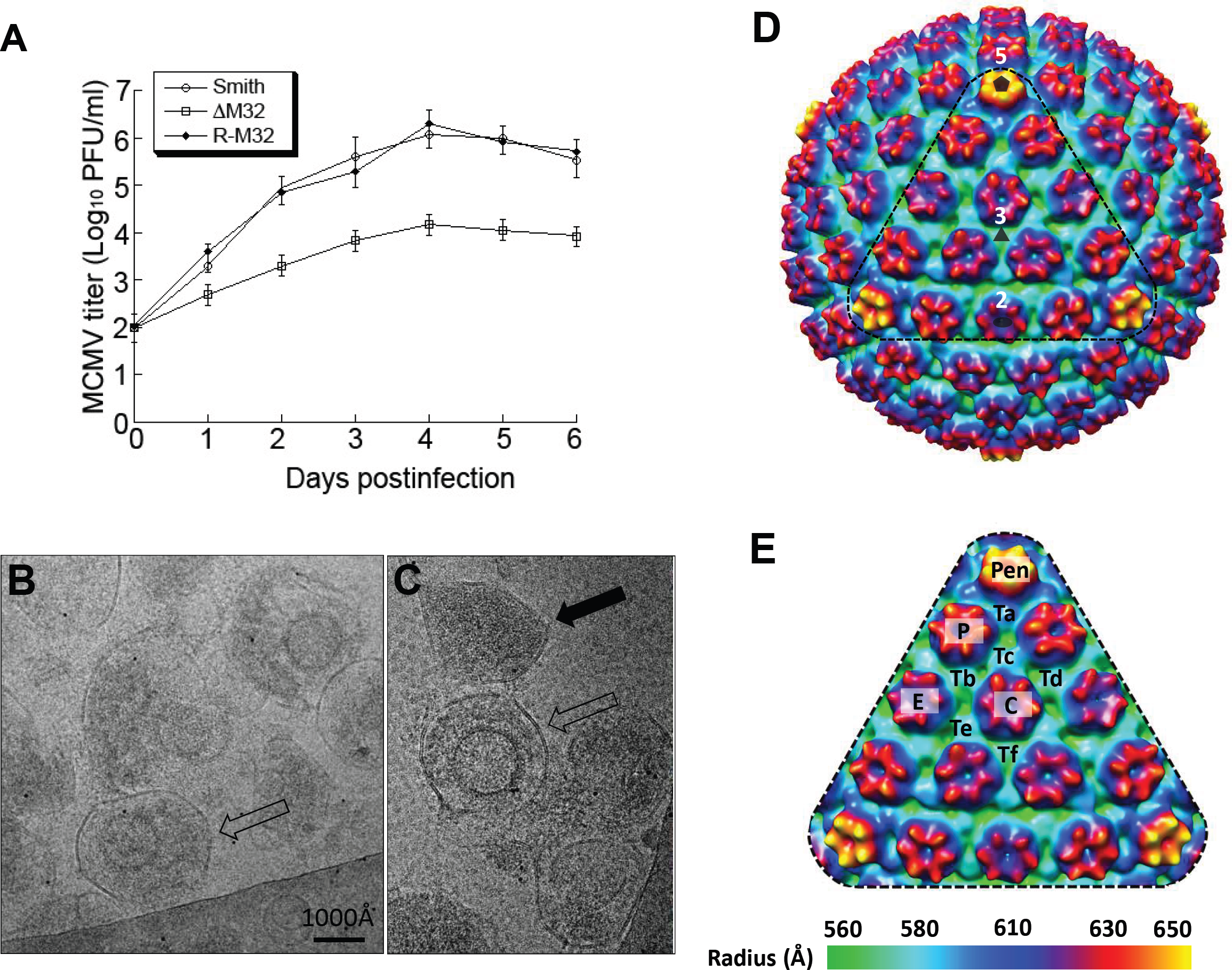
Generation and analyses of M32-deletion MCMV mutant. (**A**) Growth of the parental virus MCMV_BAC_ (Smith), M32-deletion mutant ΔM32 and rescued mutant R-M32 in NIH 3T3 cells. NIH 3T3 cells were infected with each virus at a MOI of 0.5 PFU per cell. At 0, 1, 2, 3, 4, 5, and 6 days post-infection, we harvested the cells and culture media and determined the viral titers by plaque assays on NIH 3T3 cells. The error bars indicate the standard deviations based on triplicate experiments. (**B-C**) CryoEM images of the ΔM32 MCMV mutant. Fully enveloped particle and dense body are indicated by an open and a solid black arrow, respectively. (**D**) Radially colored surface representation of cryoEM reconstruction (∼25 Å resolution) of the enveloped particles of ΔM32 MCMV, viewed along a 3-fold axis. The pentagon, triangle, and oval symbol denotes a 5-fold, 3-fold, and 2-fold axis, respectively. (**E**) Enlargement of a facet [triangle region in (D)] with pentons (Pen), hexons (C, E, P) and triplexes (Ta, Tb, Tc, Td, Te, Tf) indicated.

Though the reduced titer of ΔM32 limited the isolation of large numbers of viral particles, we still managed to purify ΔM32 viral particles. Consistent with the 100-fold reduction in its growth rate (Fig. 9A), the concentration of ΔM32 viral particles produced in the cell culture is low compared to that of the wild-type virus based on our EM analysis (Fig. 9B-C and Fig. S1A-D). Specifically, deletion of the M32 gene appeared to reduce the formation of infectious, DNA-containing virions, as it was difficult to find DNA-containing particles. Instead, most observed particles were non-infectious enveloped particles (NIEPs) and dense bodies (Fig. 9B-C), suggesting that DNA-containing particles are less stable in ΔM32. From 200 fully-enveloped cryoEM particles (an example of which is denoted by the open black arrow in Fig. 9B-C), we obtained a 3D icosahedral reconstruction of ΔM32 at ∼25 Å resolution (Fig. 9D). This resolution is sufficient to resolve pentons, hexons, and triplexes on the capsid, and as shown in the enlarged facet of the icosahedral reconstruction (Fig. 9E), the capsid of ΔM32 possesses the same molecular architecture as that of other herpesviruses (32, 33). In contrast to the ΔM32 sample, wild-type MCMV sample prepared by the same procedure demonstrated a significantly higher viral particle concentration when examined by cryoEM (Fig. S1A), and many particles are DNA-containing virions (solid black arrow in Fig. S1A). From 1,200 wild-type MCMV virion particles (*e.g.*, Fig. S1B), we obtained a ∼12 Å resolution reconstruction (Fig. S1E-G). We also obtained a reconstruction from enveloped particles without DNA (NIEPs, an example is shown in Fig. S1C), the capsid and tegument densities of which were identical to those of the virion reconstruction. Lastly, structural comparison between wild-type MCMV (∼12 Å) and the mutant ΔM32 (∼25 Å) confirmed the absence of pM32 atop triplexes (Fig. 9D-E).

## Discussion

In this study, we present the first cryoEM structures obtained from the MCMV virion and M32-deletion mutant, as well as functional data that together establish important differences concerning the structural and functional roles of pp150 in HCMV and MCMV. The attainment of an atomic model of the MCMV particle—consisting of 55 unique protein conformers of capsid and tegument proteins—is a remarkable endeavor, considering there were no atomic structures ever reported for any MCMV proteins prior to this study. Despite highly similar structures of their capsid proteins, MCMV and HCMV have distinctive capsid-tegument binding patterns: 260 “Λ”-shaped pM32nt dimers on all triplexes but Tc in each MCMV, as opposed to 320 “△”-shaped pUL32nt structures (one dimer + one monomer of pUL32nt) on all triplexes in each HCMV. Substantially more amino acids are involved in pM32-pM32 interactions in MCMV than in pUL32-pUL32 associations in HCMV, suggesting a more rigid pM32-pM32 dimer structure than pUL32-pUL32 dimer. Finally, M32-deletion mutant can be successfully generated, albeit with a growth rate reduced ∼100-fold, whereas UL32 deletion in HCMV is lethal.

The different extent of pp150-pp150 interaction and pp150 dimer rigidity in MCMV and HCMV may account for the distinctive capsid-binding patterns of pp150 in the respective viruses. The additional rigidity of pp150 dimer in MCMV may prevent the spreading of the dimer necessary to span the increased distances to adjacent SCPs surrounding the triplex Tc, presumably leading to the lack of pM32-binding in this region. The more extensive inter-pp150 interactions observed for MCMV CATC may also account for the observation that pp150 (pM32) does not exist as a monomer in MCMV, while approximately 1/3 of pp150 (pUL32) exists as a monomer in HCMV. Interestingly, HCMV pp150 is about 40% longer in sequence than its homologs in MCMV (Table S1), simian CMV, and human herpesviruses 6 and 7 (18) though a correlation has not been established between this length difference and the polymorphic difference in tegument-capsomer association due to the limited resolutions of the existing cryoEM structures of these β-herpesviruses (34, 35).

On the basis of no viral growth in cultured cells electroporated with a bacterial artificial chromosome containing the UL32-deletion HCMV genome (29), pUL32 was considered to be an essential tegument protein that can be targeted for therapeutic development against HCMV infection (14). In contrast, its counterpart in MCMV, pM32, is nonessential: M32-deletion mutant is viable *in vitro*, although defective in generating DNA-containing virions and its infectivity attenuated by 100-fold (Fig. 9A). Evidence shows that “SCP-deficient” HCMV viral particles have decreased viral yield (10,000-fold) compared to that of wild-type virus (36). As mentioned previously, SCP structures in MCMV and HCMV are highly conserved. Thus, it is worth considering SCP as a drug target while rationally designing novel drugs against MCMV infections. While our results establish the structural basis of using MCMV as a model for HCMV pathogenesis and therapeutic studies when targeting capsid proteins such as SCP, caution is warranted when targeting tegument protein pp150 due to its different structural organization and functional roles in MCMV and HCMV reported in this study.

## Method

### Isolation of MCMV virions

Mouse NIH3T3 cells were cultured in Dulbecco’s Modified Eagle Medium (DMEM) plus 10% fetal bovine serum (FBS). Twenty flasks (175 cm^2^ each) of cells were grown to 90% confluence and then infected with MCMV Smith strain at a multiplicity of infection (MOI) of 0.1. At 6 days post infection, when half of the cells were lysed, the media was collected and centrifuged at 10,000g for 15 min to remove cell debris. The clarified supernatant was then collected and centrifuged at 80,000g for 1 hr to pellet MCMV virions. Pellets were resuspended in a total volume of 2 ml phosphate buffered saline (PBS, pH 7.4) and loaded on a 15%-50% (w/v) sucrose density gradient and centrifuged at 100,000g for 1 hr. We usually observe three light-scattering bands—top, middle, and bottom – containing mainly noninfectious enveloped particles (NIEPs), virions, and dense bodies, respectively. The middle band (virions) was collected and diluted in PBS to a total volume of 13 ml. Virion particles were pelleted again at 80,000g for 1 hr and resuspended in 30 μl PBS for cryoEM sample preparation.

### CryoEM data acquisition

Purified intact MCMV virions were mixed with NP-40 detergent at 1% final concentration to partially solubilize the viral envelope. Immediately after, aliquots of 2.5 μl of this treated sample were applied to 200-mesh Quantifoil R2/1 grids, blotted with filter paper, and plunge-frozen in liquid ethane. CryoEM images were collected at liquid nitrogen temperature in an FEI Titan Krios cryo electron microscope operated at 300 kV with parallel illumination. Images were recorded on Kodak SO163 films with a dosage of ∼25 e^−^/Å^2^ at 47,000× nominal magnification. A total of 2,200 films were recorded and digitized using Nikon Super CoolScan 9000 ED scanners at 6.35 μm per pixel (corresponding to 1.351 Å per pixel at the sample level).

### CryoEM data processing and structure determination

Defocus values of all micrographs were determined with *CTFFIND3* (37) to be in the range of −1μm to −3μm. Particles were picked with *Ethan* (38) and then manually screened with the boxer program in *EMAN* (39) to keep only well-separated and artifact-free particles. A total of 58,254 particle images were boxed out from the micrographs with *EMAN*. The original particle images were binned 8×, 4×, or 2× stepwise to speed up data processing. Icosahedral refinement and reconstruction were carried out with the common line-based *IMIRS* package (40, 41) and GPU-implemented reconstruction program *eLite3D* (42), respectively. The final capsid reconstruction was obtained by averaging 47,982 particles.

#### Sub-particle reconstructions for 5-fold, 3-fold, and 2-fold axis regions

Due to the large size of the MCMV capsid (over 1,300 Å in diameter), resolution of our initial icosahedral reconstruction was limited to 5 Å, likely due to slight particle deformation and defocus gradient across the depth of the sample (12, 19). To obtain higher resolution structures for reliable atomic model building, we applied a localized reconstruction strategy (12, 43) to reconstruct subareas surrounding the 2-fold, 3-fold, and 5-fold axes of the icosahedral MCMV capsid. The 47,982 high-quality particle images selected from IMIRS refinement were binned 4x and reprocessed using *Relion* (44) with icosahedral symmetry applied. Using the script downloaded from www.opic.ox.ac.uk/localrec (43), positions of sub-particles in the original particle images (*i.e.*, without binning) were calculated with a distance of 576.8 Å from the center of the viral particle. A total of 575,784; 959,640; and 1,439,460 sub-particles in 400 x 400 pixels were then boxed out for the 5-fold, 3-fold, and 2-fold axis, respectively. Localized reconstruction of these sub-particles were iteratively refined in *Relion*, reaching a final estimated resolution of 3.8, 3.6, and 3.8 Å for the 5-fold axis, 3-fold axis, and 2-fold axis sub-particle maps, respectively, based on the 0.143 FSC criterion (45).

#### Averaging density maps of the triplex regions

To improve the quality of the pM32 dimer density, we obtained sub-particle reconstructions of triplexes Ta, Tb, Td, and Te regions and performed averaging using the Tb, Td, and Te sub-particle reconstructions as per the work flow outlined in Figure S3. Briefly, sub-particles centered on triplexes Ta, Tb, Td, and Te were extracted in boxes of 150 × 150 pixels and reconstructed separately using the same procedure for sub-particle reconstructions of the 5-fold, 3-fold, and 2-fold axis regions described above. These sub-particle reconstructions show that pM32 dimers in the Tb, Td, and Te sub-particle reconstructions are highly similar to each other, though less so with the Ta sub-particle reconstruction. We therefore performed sub-particle refinements for Tb, Td, and Te regions and averaged their resulting sub-particle reconstructions to further improve the signal-to-noise ratio. Density maps were first manually and then computationally aligned by the “*fit in map*” function in *Chimera* (46). An averaged map for the triplex region was obtained by executing the “*vop add*” command in *Chimera* on the above aligned density maps.

#### Sub-particle classification for the triplex Tc region

To verify whether the lack of pM32 binding on triplex Tc is due to icosahedral symmetrization, we performed sub-particle 3D classification for the triplex Tc region, following the work flow outlined in Figure S4. Sub-particles were extracted in a box of 150 × 150 pixels centered on triplex Tc and processed by 3D classification with ten classes requested. All resulting 3D classes possessed triplex density but lacked pM32 density.

#### Sub-particle classification and refinement for triplex Tf

Due to the imposition of icosahedral symmetry during 3D reconstruction, the tegument protein density associated with triplex Tf was smeared (Fig. 1B). To overcome this problem, we extracted sub-particles from the Tf region in a box of 150 × 150 pixels, expanded them with C3 symmetry, and performed 3D classification and refinement with C1 symmetry, following the work flow outlined in Figure S5. We obtained three similar 3D structures related to each other by 120 degrees of rotation (last row in Fig. S5), the second reconstruction (#2) had the highest resolution (6.8 Å) with the orientation of the triplex distinguishable. Refining this structure revealed a pM32 dimer.

### Atomic model building and refinement

Each asymmetric unit of the T=16 icosahedral reconstruction of the MCMV particle contains 55 unique copies of protein subunits: 16 MCPs, 16 SCPs, 15 triplex subunits (excluding triplex Tf, the resolution of which at 6.8 Å is insufficient for atomic model building), and 8 pM32s. We utilized the *SWISS-MODEL* server (47) to generate homology models of penton MCP, hexon MCP, SCP, Tri1, Tri2A, and Tri2B with the corresponding subunit conformers in the atomic model of HCMV (14) as templates. These initial models were docked into the sub-particle reconstructions (sharpened with a B factor of −150 Å^2^for the 5-fold axis map, −160 Å^2^for the 3-fold axis map, and −160 Å^2^ for the 2-fold axis map) and then adjusted manually in *Coot* (48). All models were then iteratively improved by *Phenix* real space refinement (49) and manual readjustment in *Coot*. Eventually, all the atomic models built based on high-resolution sub-particle reconstructions were assembled together and refined against the icosahedral reconstruction with *Phenix* to correct for inter-molecular clashes.

Unlike for the capsid proteins, the atomic model for pM32nt was built *de novo*. The resolution of densities corresponding to pM32nt was poorer than that of capsid proteins even in the improved sub-particle reconstructions. Thus, the map obtained by averaging triplexes Tb, Td, and Te regions as previously mentioned was also used to facilitate atomic model building for pM32nt. Secondary structures predicted by *Phyre2* server (50) were used to guide backbone tracing using the *Baton_build utility* in *Coot*. Observable main chain residue bumps in the density informed Cα placement, and registration was accomplished using distinguishable side chain densities. Iterative model refinement was performed as for the capsid proteins described above.

### Generation of M32-deletion mutant and wild-type MCMV

We used a previously reported bacterial artificial chromosome (BAC)-based clone of MCMV Smith strain (MCMV_BAC_), maintained as a BAC-based plasmid in *E. coli*, to produce infectious progeny in mouse NIH3T3 cells (28-30). As reported, this progeny retained wild-type growth characteristics *in vitro*, as previously shown (28-30).

Mutant ΔM32, which contained a deletion of the entire coding sequence of M32, was derived from MCMV_BAC_ using a two-step mutagenesis protocol (29, 31). In the first step, we inserted at the M32 sequence with a cassette (tet/str) containing the tetracycline resistance gene tetA and rpsL gene conferring streptomycin susceptibility, following the mutagenesis procedures as described previously (29, 31). The bacteria harboring the mutant BAC constructs were electroporated with the PCR-amplified tet/str cassette. Successful insertion of the tet/str cassette was screened by selecting for bacterial colonies resistant to tetracycline. In the second step, the tet/str cassette was targeted for deletion. The resulting mutant, which only contained the deletion of M32 ORF sequence, was streptomycin-resistant and, therefore, was easily selected in the presence of the antibiotics (29, 31). Rescued mutant R-M32 was generated from ΔM32, following the experimental procedures for construction of rescued viruses as described previously (29, 31). The M32 regions in the mutant and the rescued virus were analyzed by restriction digestion profile and sequencing analyses.

Virus growth analyses were carried out by infecting NIH 3T3 cells (n =1×10^6^) with viruses (MOI = 0.5-1) (51, 52). The cells and medium were harvested at different time points postinfection. Viral stocks were prepared and used to infect NIH3T3 cells, followed by agar overlay. Viral titers were determined by counting the number of plaques 5-7 days after infection (51, 52). The values obtained were averages from three independent experiments.

For cryoEM, we used a sucrose density gradient to purify viral particles from the supernatant of culture media of wild-type or ΔM32-infected NIH3T3 cells as previously described (27). The sucrose gradient fraction with the most viral particles from the wild-type MCMV prep and the corresponding fraction from the ΔM32 prep were collected for cryoEM imaging. We suspended a 3 μl aliquot of each of these samples to a holey carbon-coated cryoEM grid, which was blotted and immediately plunge-frozen so that viral particles were suspended within vitreous ice across holes of the holey carbon (53). Low-dose (∼20 e^-^/Å^2^) cryoEM images were recorded on a Gatan 16-megapixel CCD camera in a Titan Krios cryo electron microscope operated at 300 kV at a magnification of 79,176× with Leginon (54). Consistent with the 100-fold reduction in its growth rate, the concentration of ΔM32 viral particles produced in cell culture is low when compared to that of the wild-type virus. Because it was hard to find viral particle-containing sample regions for imaging, many imaging sessions were painstakingly carried out to obtain just 500 micrographs for ΔM32. (On average, each micrograph only contained one viral particle.) We selected 200 fully-enveloped cryoEM particles (see an example pointed by the open black arrow in Fig. 9B-C) to reconstruct a 3D structure of ΔM32 at ∼25 Å resolution with the *IMIRS* package (40, 41). In contrast to the ΔM32 sample, wild-type MCMV preparation obtained from the same procedure had a significantly higher viral particle concentration. 3D reconstructions obtained from wild-type virions and NIEPs with *IMIRS* were thus at higher resolutions than ΔM32 due to the larger number of particles used.

## Data deposition

CryoEM maps and atomic models are deposited in the Electron Microscopy Data Bank (EMDB) and the RCSB Protein Data Bank (PDB), respectively. They include the cryoEM density maps of the MCMV capsid, sub-particle reconstructions at 5-fold, 3-fold, and 2-fold axes (accession code EMD-XXXXX, XXXXX, XXXXX, XXXXX, respectively) and a single coordinate file containing 55 atomic models (PDB accession code XXXX).

## Acknowledgments

We thank Xing Zhang for assistance in cryoEM, Sean Umamoto, Hao Gong, Xiaohong Jiang and Eric Huynh for M32-deletion mutant virus work. This project is supported in part by grants from the National Institutes of Health (DE025567, AI069015 and GM071940 to ZHZ) and (DE023935 and DE025462 to FL) and Guangdong Innovative and Entrepreneurial Research Team Program (No. 2014ZT05S136 to FL). We acknowledge the use of cryoEM facilities at the Electron Imaging Center for Nanomachines supported in part by NIH (1S10RR23057) and NSF (DMR-1548924). WL acknowledges support from the China Scholarship Council.

## Supplementary information

**Table. S1.** Comparison of capsid and pp150 in MCMV and in HCMV.

| Protein names | Size (a.a.) |  | Sequence (%) |  |
| --- | --- | --- | --- | --- |
|  | HCMV | MCMV | Identity | Similarity |
| MCP | 1370 | 1353 | 56.1 | 72.2 |
| SCP | 75 | 98 | 34.4 | 65.6 |
| Tri1 | 290 | 294 | 36.3 | 57.6 |
| Tri2 | 306 | 311 | 54.4 | 73.4 |
| pUL32/pM32 | 1046 | 718 | 27.9 | 43.0 |

## Supplementary figure legends

**FIG. S1.**
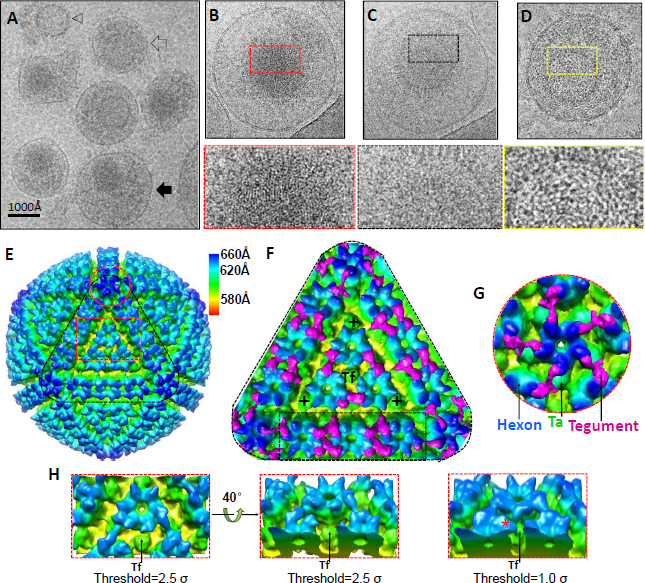
CryoEM imaging of wild-type MCMV (without detergent treatment) and 3D reconstruction of its icosahedral capsid. (**A**) A cryoEM micrograph of wild-type MCMV, containing virions (solid black arrow), a non-infectious enveloped particle (NIEP, open black arrow), and a naked capsid (black arrowhead). (**B-D**) Representative enveloped MCMV particles with DNA (B; *i.e.*, virion), without DNA (C; *i.e.*, NIEP), and with leaked out DNA (D). A characteristic “finger-print” pattern of closely packed viral DNA is visible in (B), not in the NIEP (C). Presumably, a portion of DNA can leak out from a structurally-compromised capsid, a characteristic fingerprint pattern of closely packed DNA becomes loose and “spaghetti-like”, spreading out into space beyond the capsid within the virion (D). These DNA features are shown more clearly in the zoom-in views of the upper portion of the capsid at the bottom of each panel. (**E**) Radially colored surface representation of the 3D icosahedral reconstruction of the wild-type MCMV virion at ∼12 Å resolution, viewed along a 3-fold axis. (**F-G**) Enlargement of a facet (F) [the triangle region in (E)] and penton (G) area [the red circle region in (E)] showing the “Λ”-shaped tegument proteins (magenta) connecting triplexes, hexons and pentons. (**H**) The boxed area of (E) observed from two different views and at two different density thresholds. Tegument densities (one marked by an asterisk) are visible and are associated with triplex Tf in the right panel when the map is rendered at a threshold of 1.0 standard deviation (σ).

**FIG. S2.**
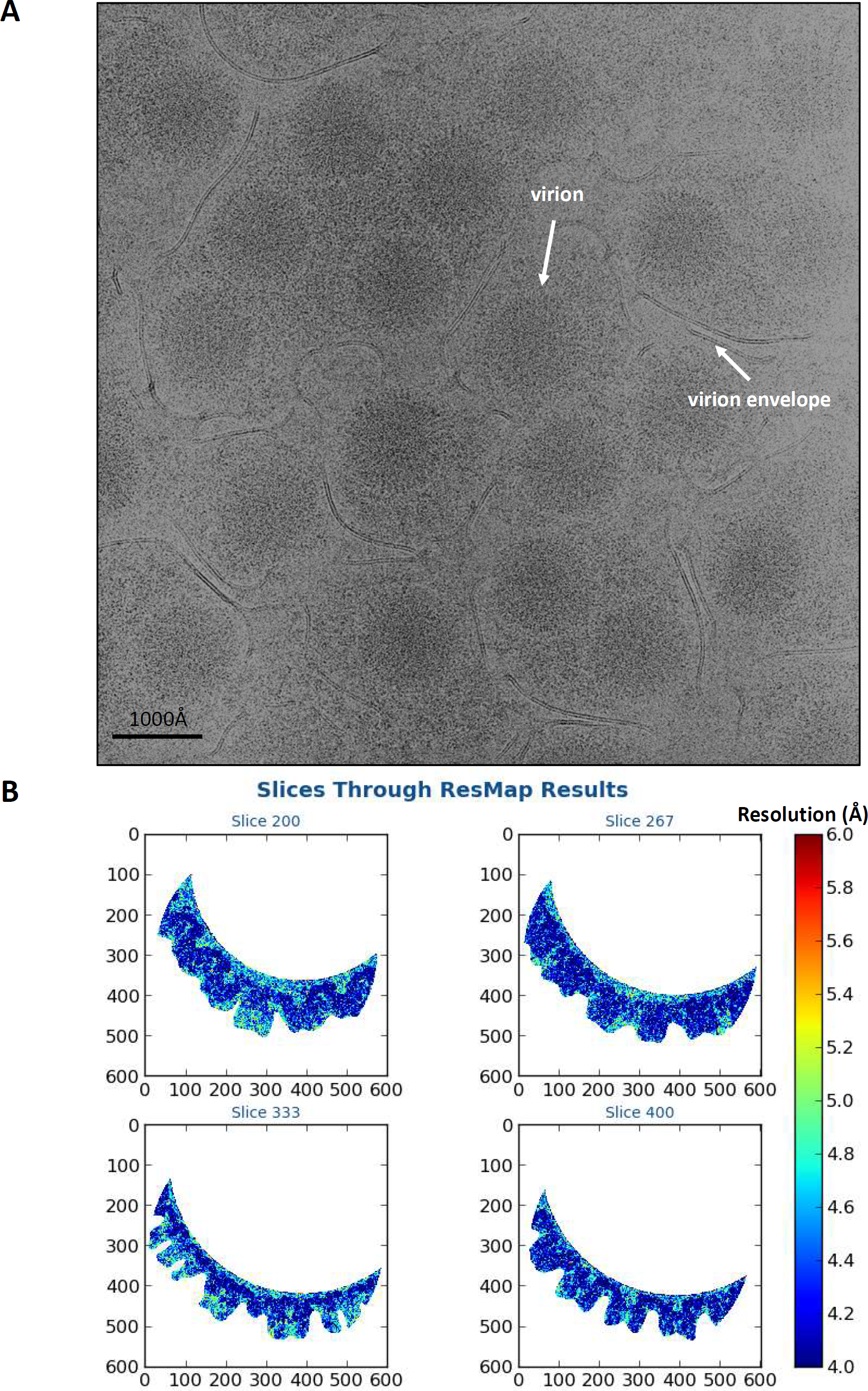
CryoEM imaging of MCMV particles with mild detergent treatment and local resolution assessment. (**A**) Image recorded on photographic film shows a high viral particle concentration. (**B**) Local resolution heat maps of the density slices through an asymmetric unit, obtained with *ResMap* (A. Kucukelbir, F. J. Sigworth, and H. D. Tagare, Nat Methods 11:63-65, 2013. doi:10.1038/nmeth.2727). Color scheme for local resolutions is shown in the color bar.

**FIG. S3.**
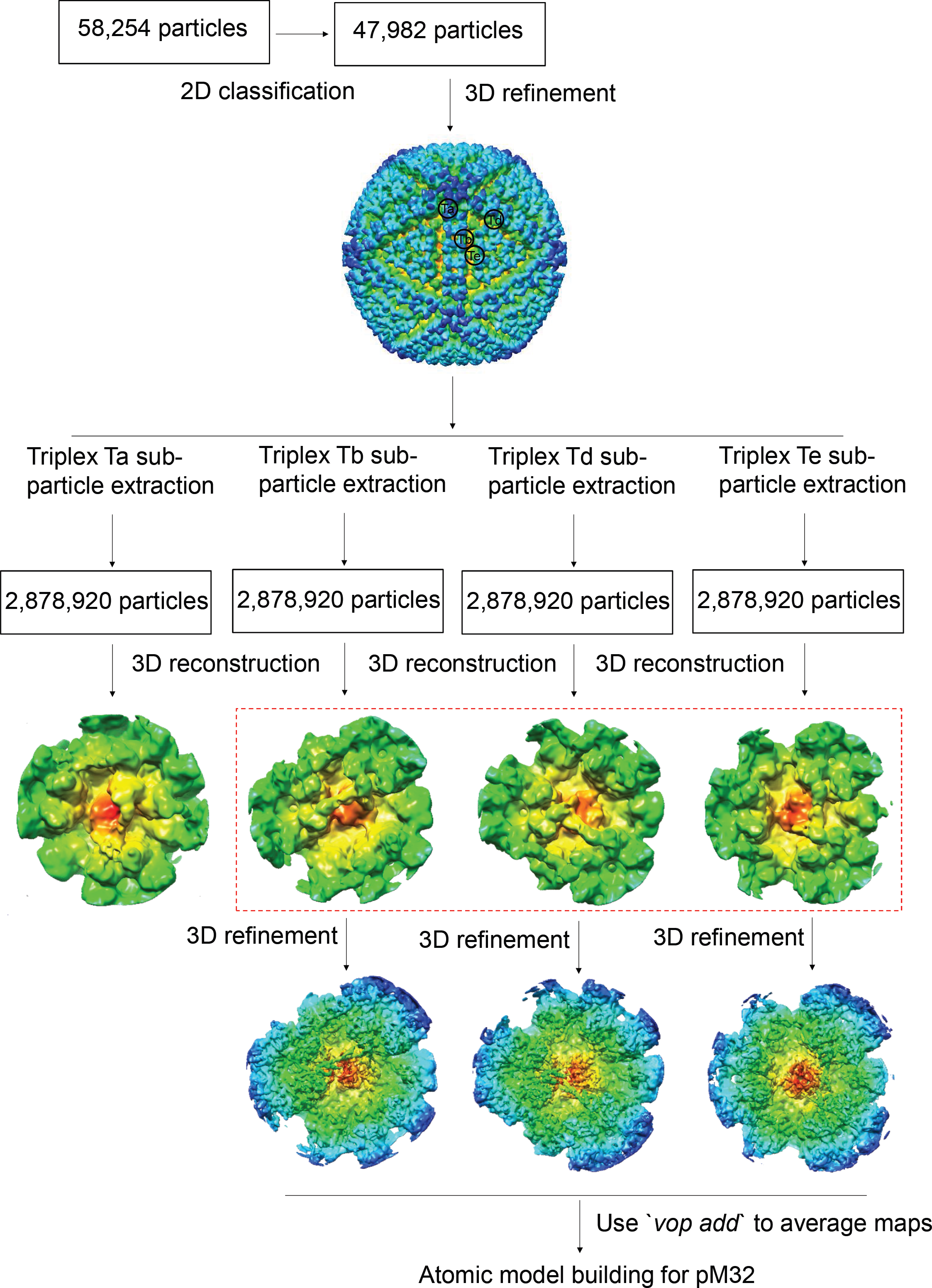
Work flow for sub-particle reconstruction of triplex (Ta, Tb, Td, and Te) regions to improve pM32 structure. Viral particles were sorted out and subjected to 2D classification in *Relion* to select only virion particles for icosahedral reconstruction. Sub-particles from triplex regions (Ta, Tb, Td, and Te) were extracted and reconstructed separately. Sub-particles from triplex Tb, Td, and Te regions were further refined and averaged by ‘*vop*’ tool in *Chimera* to boost the signal-to-noise ratio of the density map. pM32 densities in the resulting averaged map was used for atomic model building.

**FIG. S4.**
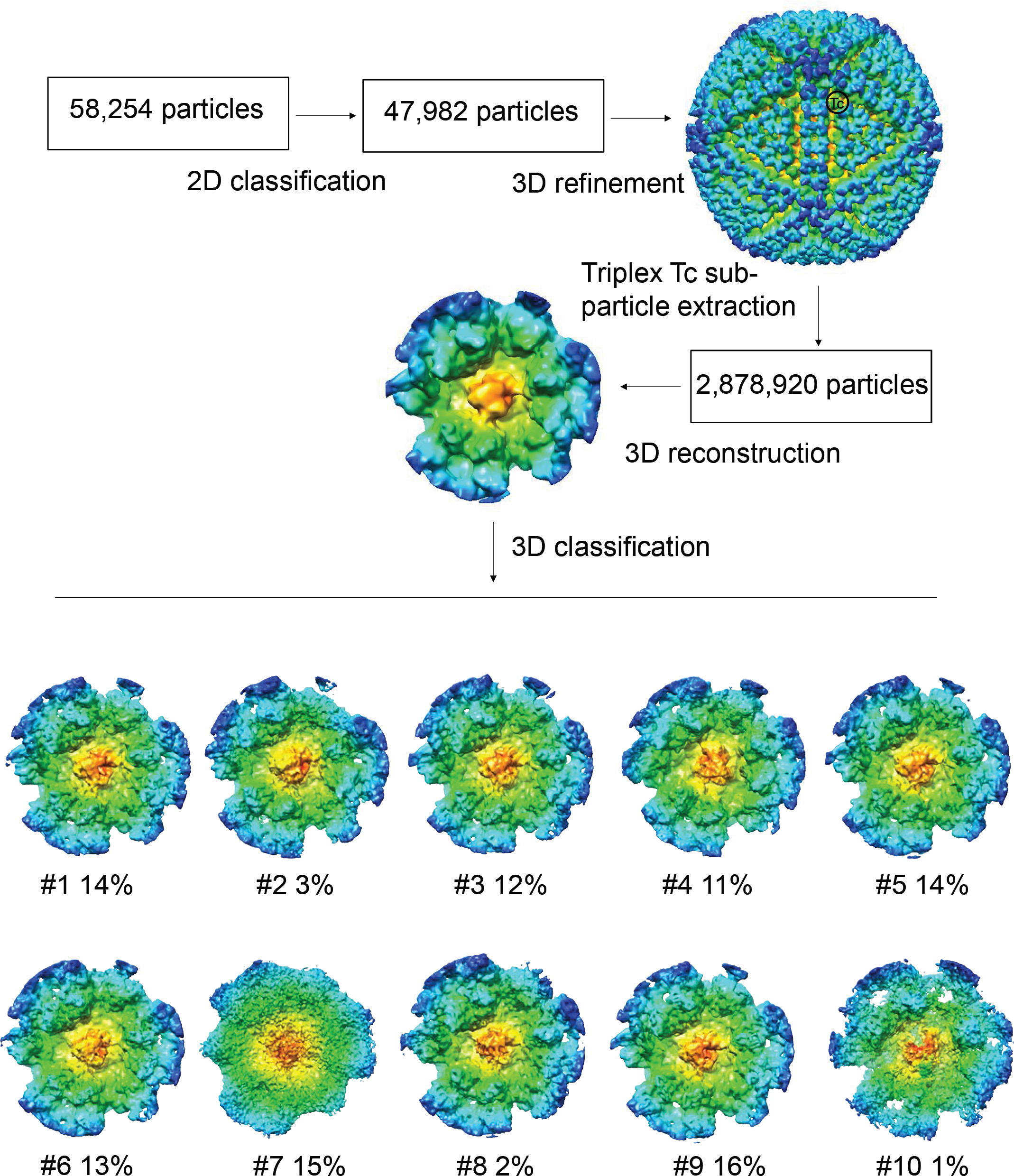
Work flow for sub-particle reconstruction and 3D classification for triplex Tc region. Sub-particles from triplex Tc region were extracted and processed by 3D classification to confirm the absence of pM32 on triplex Tc.

**FIG. S5.**
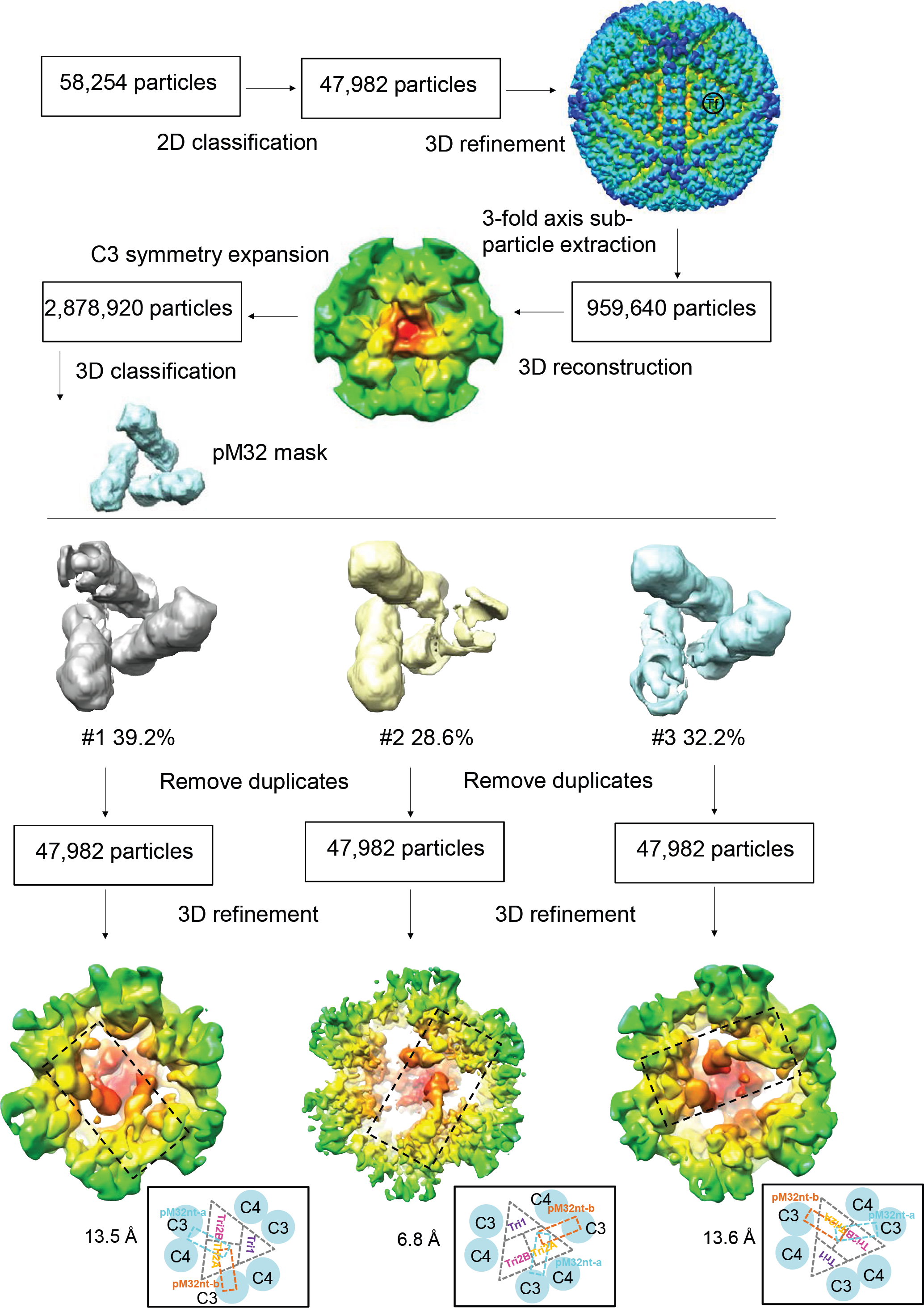
Work flow for sub-particle reconstruction and 3D classification for triplex Tf region. Sub-particles containing tegument contributions above triplex Tf were extracted with a 3-fold symmetric pp150 mask and subjected to 3D classification and refinement. All the resulting reconstructions contain only two pM32 subunits.

**FIG. S6.**
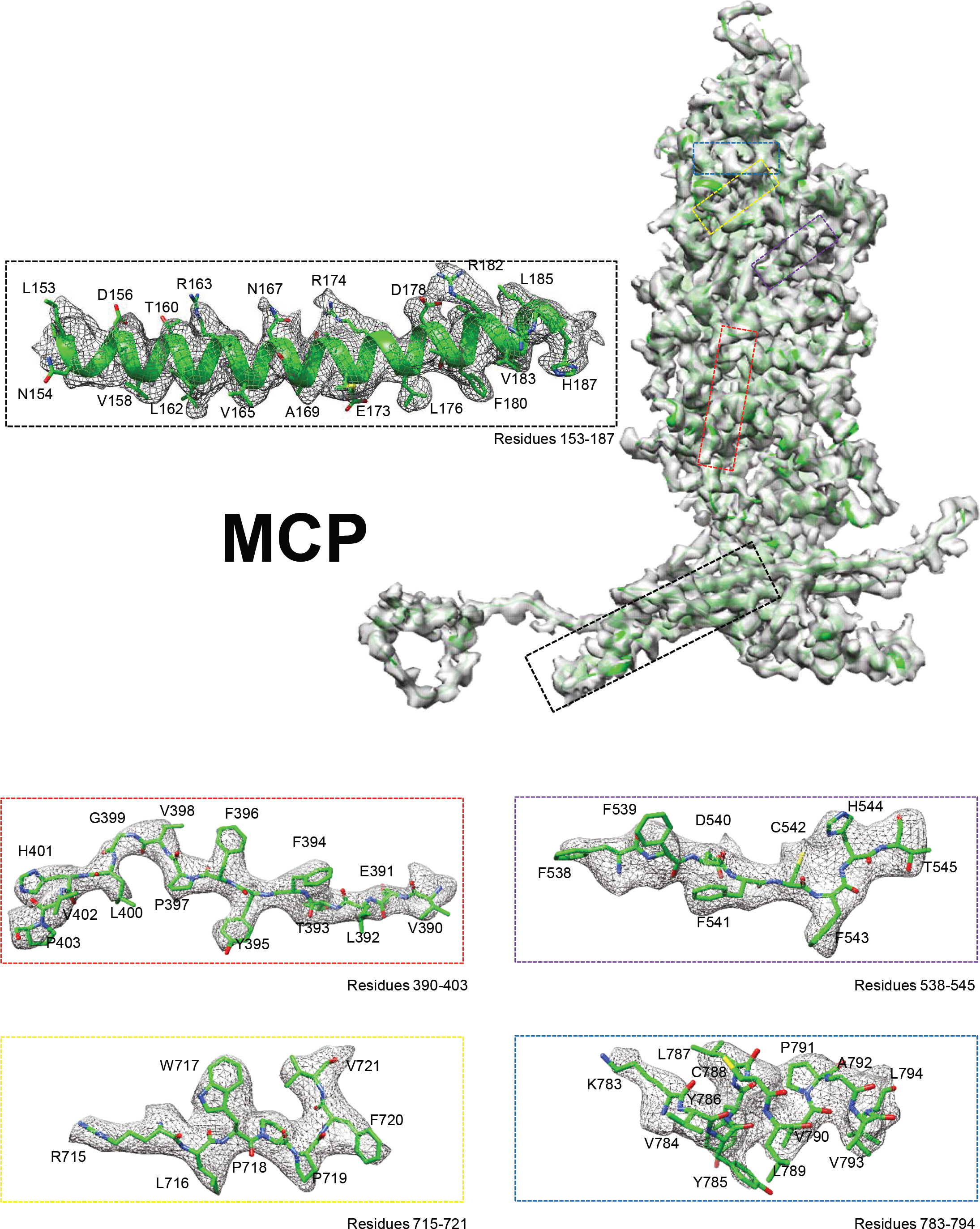
Density map and atomic model of a hexon MCP monomer. The density map (gray) of a hexon MCP segmented out from the 3-fold axis sub-particle reconstruction (3.6 Å) is superposed with its atomic model (ribbon). Boxed regions are enlarged with density shown as gray mesh and atomic models as ribbon/sticks in the boxes with corresponding color edges.

**FIG. S7.**
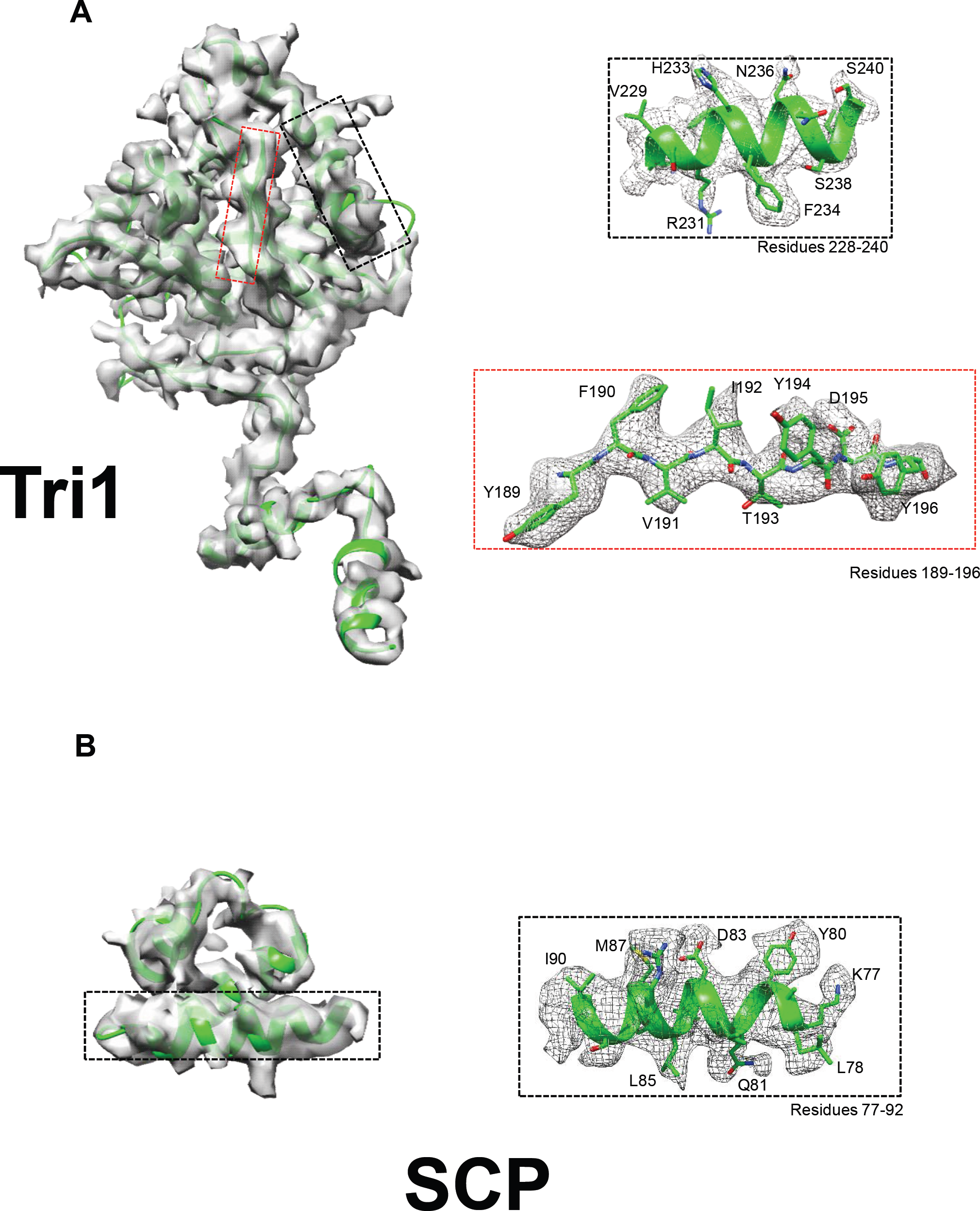
Density maps and atomic models of a Tri1 and an SCP monomer. (**A-B**) The density maps (gray) of a Tri1 (A) and an SCP (B) segmented out from the 3-fold axis sub-particle reconstruction (3.6 Å) are superposed with their atomic models (ribbon). Boxed regions are enlarged with density shown as gray mesh and atomic models as ribbon/sticks in the boxes with corresponding color edges.

**FIG. S8.**
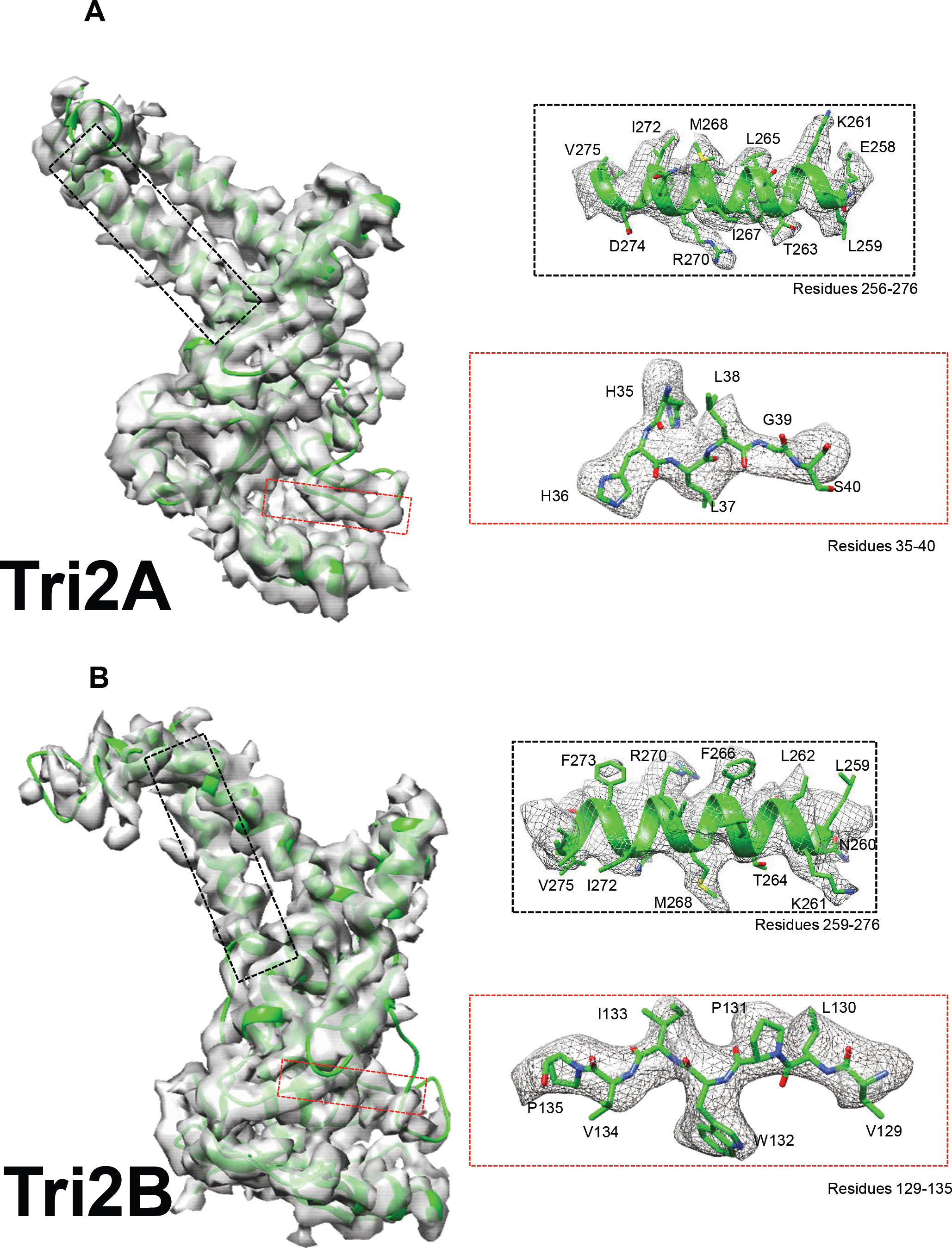
Density maps and atomic models of a Tri2A and a Tri2B monomer. (**A-B**) The density maps (gray) of a Tri2A (A) and a Tri2B (B) segmented out from the 3-fold axis sub-particle reconstruction (3.6 Å) are superposed with their atomic models (ribbon). Boxed regions are enlarged with density shown as gray mesh and atomic models as ribbon/sticks in the boxes with corresponding color edges.

**FIG. S9.**
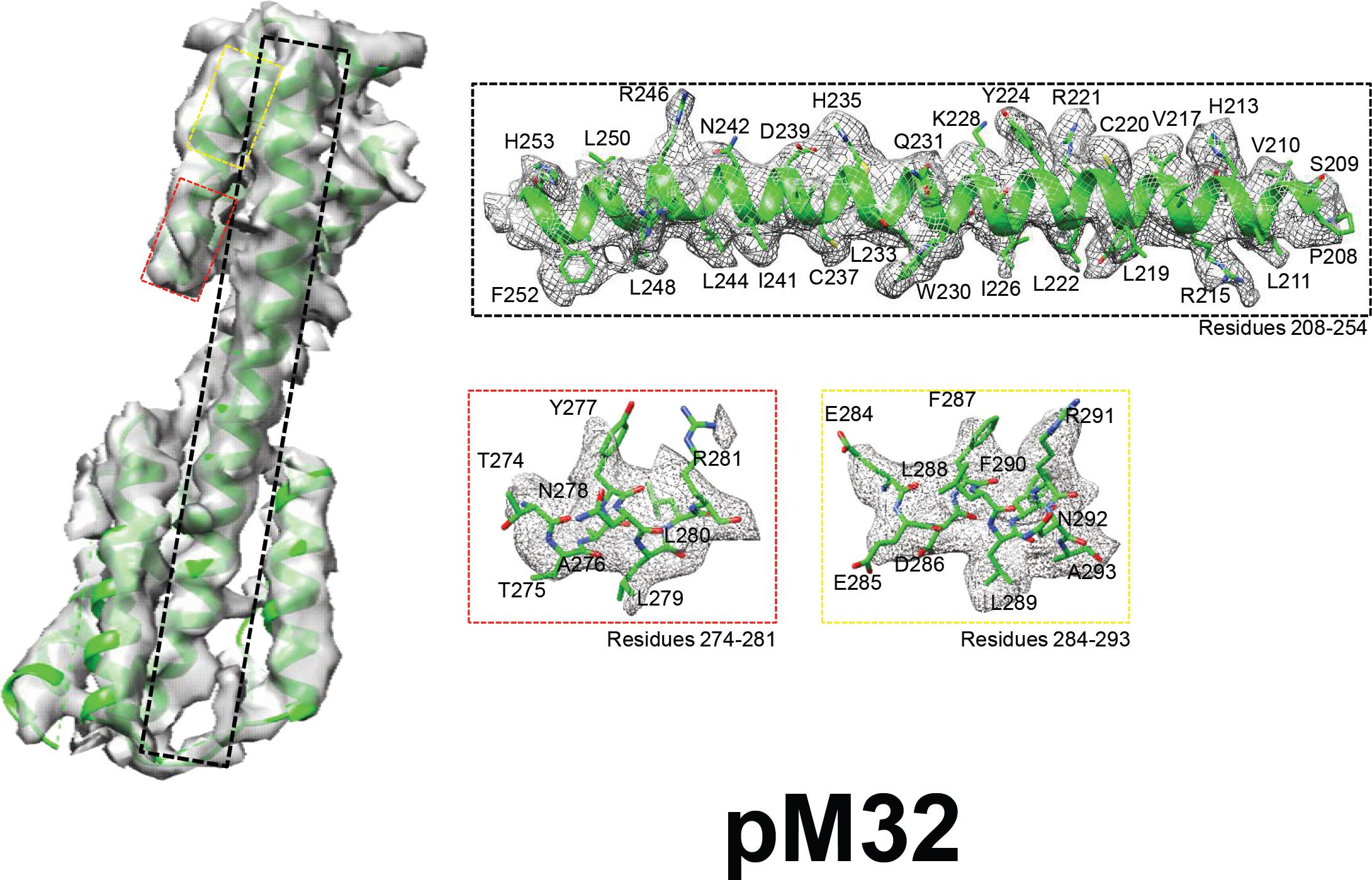
Density maps and atomic models of a pM32nt monomer. The density map (gray) of a pM32 segmented out from the 3-fold axis sub-particle reconstruction (3.6 Å) is superposed with its atomic model (ribbon). Boxed regions are enlarged with density shown as gray mesh and atomic models as ribbon/sticks in the boxes with corresponding color edges.

**FIG. S10.**
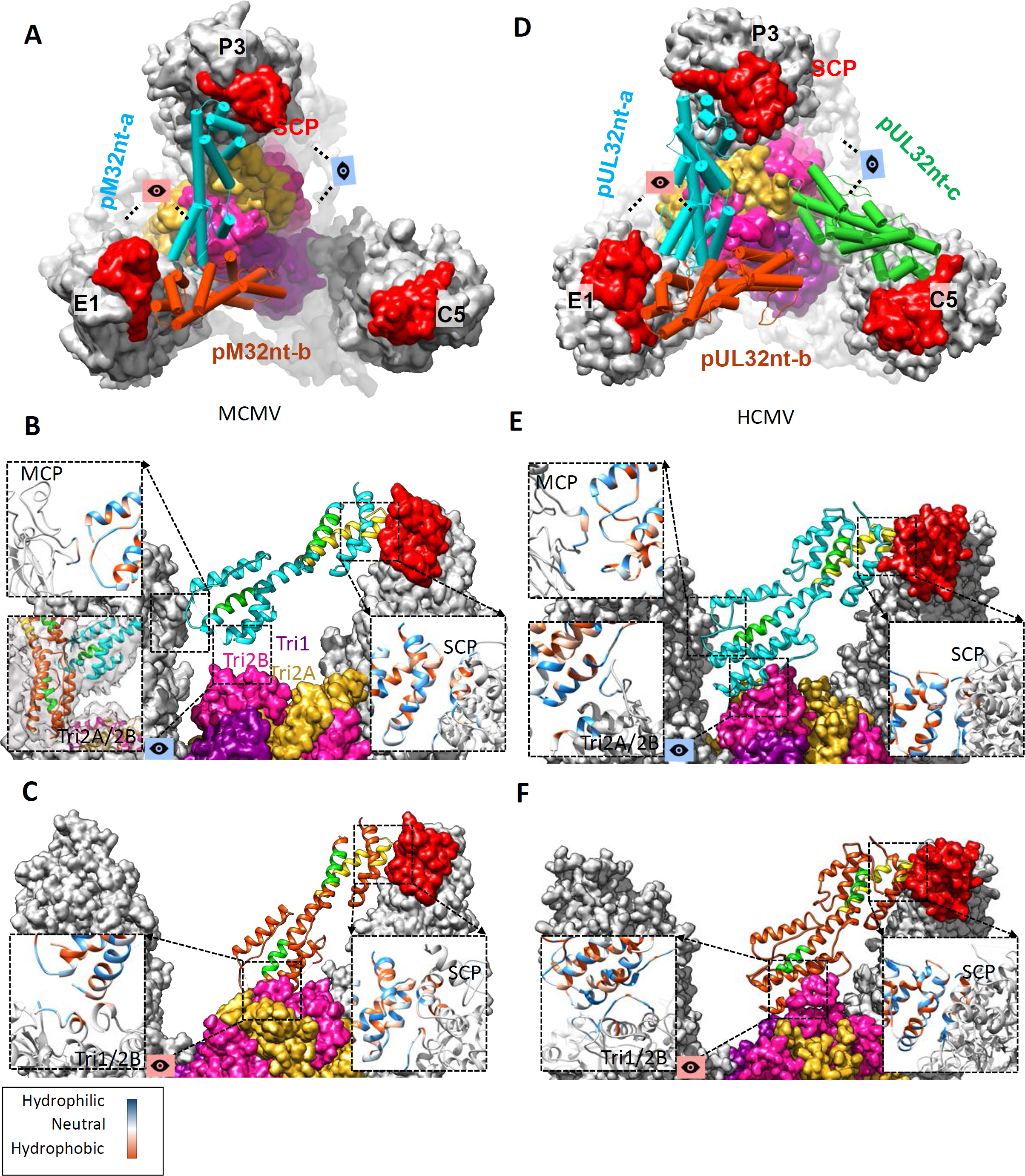
The securing of tegument protein to capsid. (**A**) Top view (down a local 3-fold axis) of triplex Td region showing a pM32 dimer—pM32nt-a (cyan) and pM32nt-b (orange red)—bound to the triplex and extending towards the SCPs atop nearby MCPs in MCMV. (**B-C**) Side views of the structure in (A), showing how pM32nt-a (B) and pM32nt-b (C) interact with capsid proteins. (**D-F**) Corresponding views in HCMV.

**FIG. S11.**
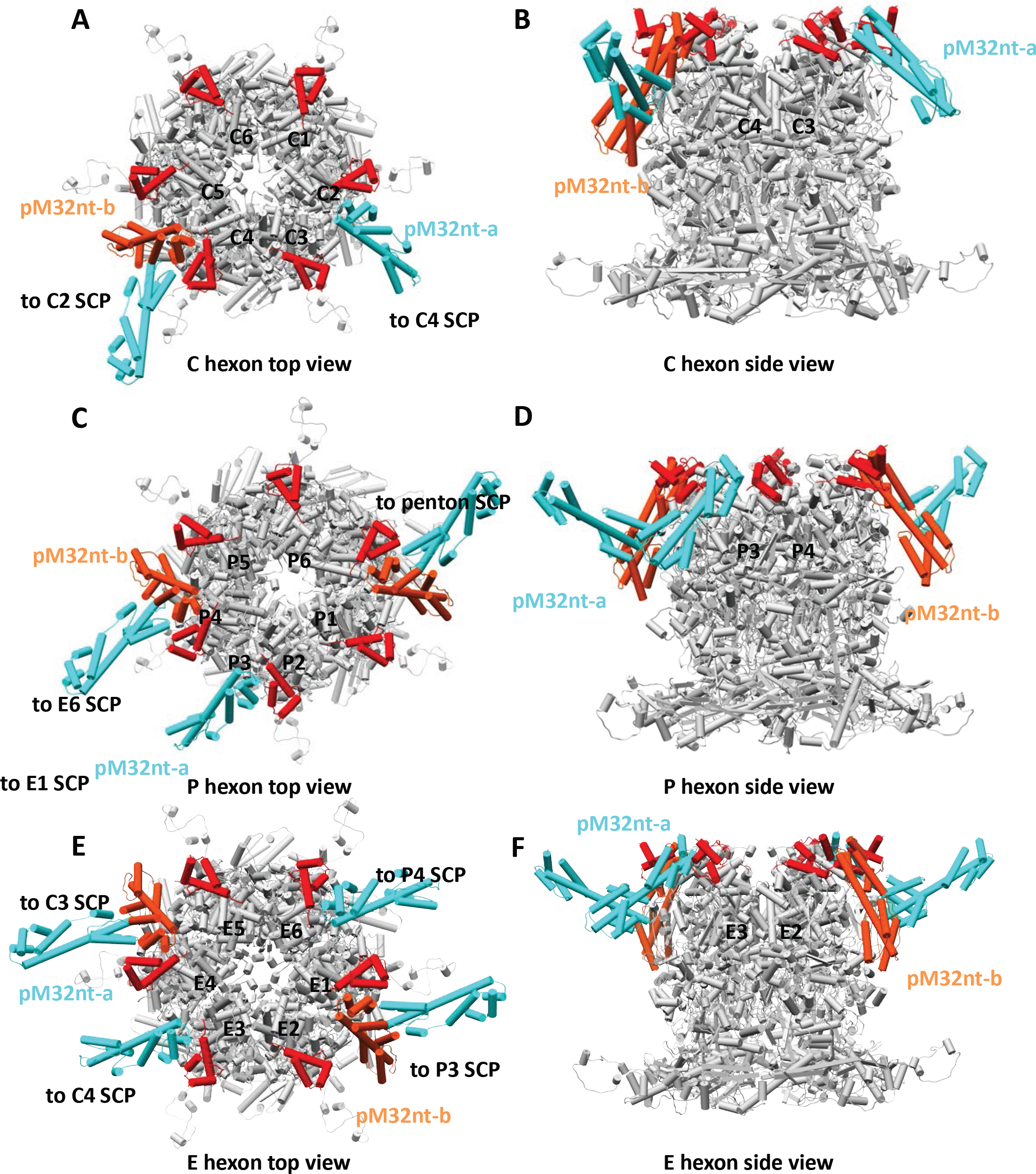
Stabilization of hexon and penton capsomers by pM32nt. (**A**) Top view of a C capsomer and its interacting pM32nt subunits, showing C capsomer is stabilized by three copies of pM32nt. (**B**) Side view of (A). (**C**) Top view of a P capsomer and its interacting pM32nt subunits, showing P capsomer is stabilized by five copies of pM32nt. (**D**) Side view of (C). (**E**) Top view of an E capsomer and its interacting pM32nt subunits, showing E capsomer is stabilized by six copies of pM32nt. (**F**) Side view of (E).

**FIG. S12.**
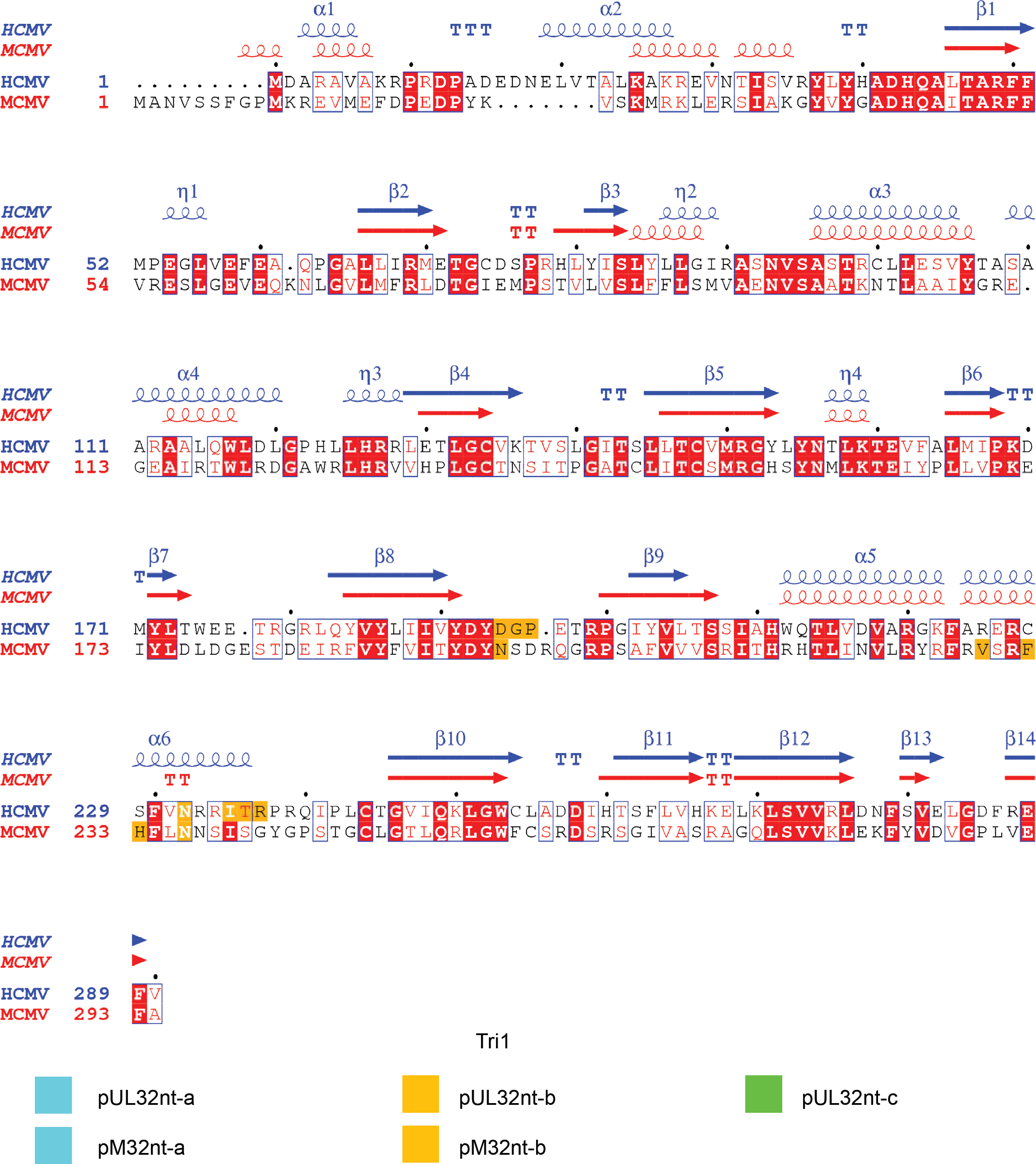
Secondary structure and sequence alignment for Tri1 in MCMV and HCMV. Schematic representations of the amino acid sequence and secondary structure alignment for Tri1 in HCMV and MCMV analyzed and displayed by *ESPript 3.0* (X. Robert and P. Gouet, Nucleic Acids Res 42:W320-W324, 2014, doi: 10.1093/nar/gku316). Arrow and spiral represent β-sheet and α-helix, respectively. Residues in Tri1 interacting with pUL32-a/pM32-a (cyan), pUL32-b/pM32-b (orange red), and pUL32-c (green, only in HCMV) are colored.

**FIG. S13.**
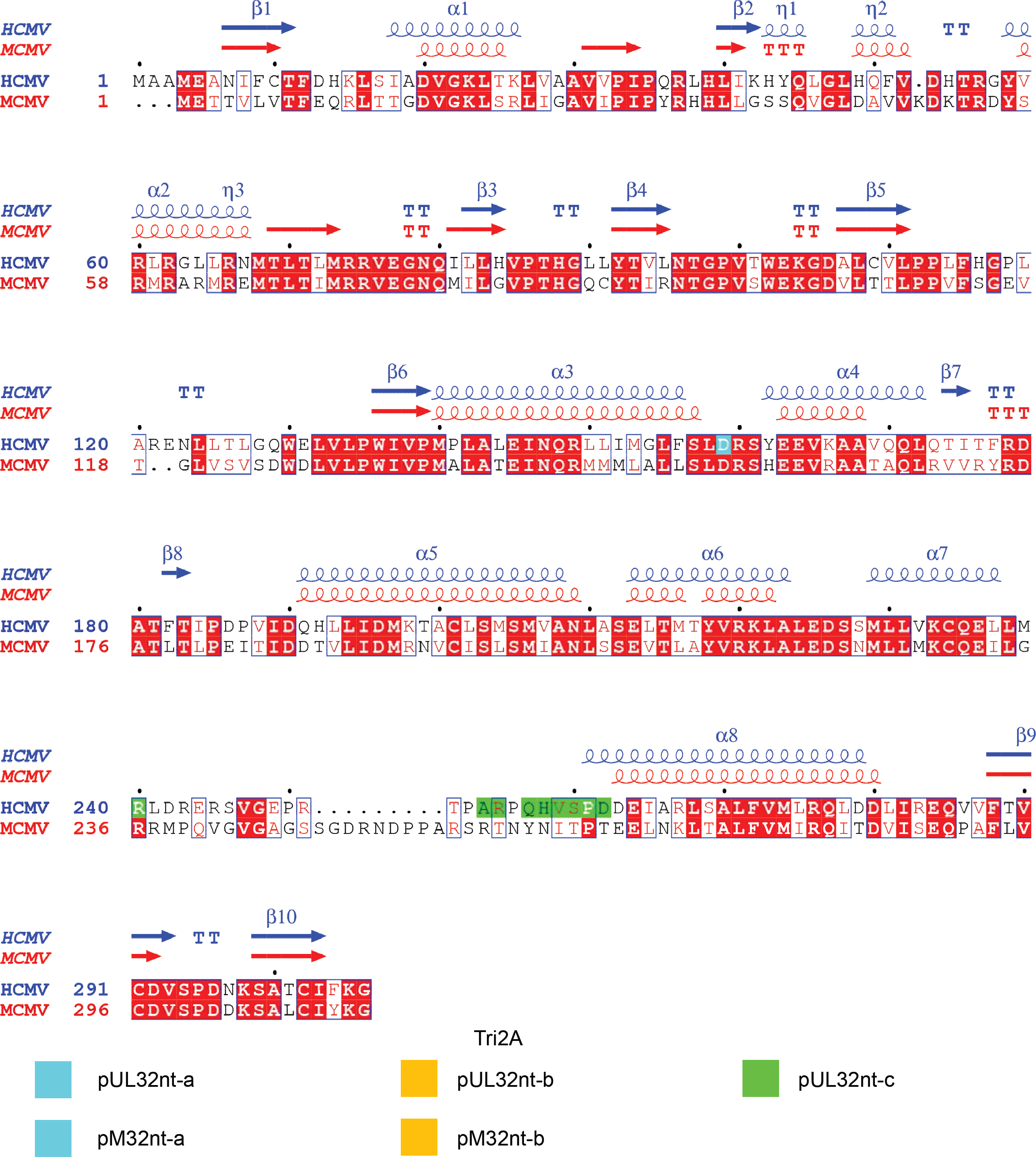
Secondary structure and sequence alignment for Tri2A in MCMV and HCMV. Schematic representations of the amino acid sequence and secondary structure alignment for Tri2A in HCMV and MCMV analyzed and displayed by *ESPript 3.0*. Arrow and spiral represent β-sheet and α-helix, respectively. Residues in Tri2A interacting with pUL32-a/pM32-a (cyan), pUL32-b/pM32-b (orange red), and pUL32-c (green, only in HCMV) are colored.

**FIG. S14.**
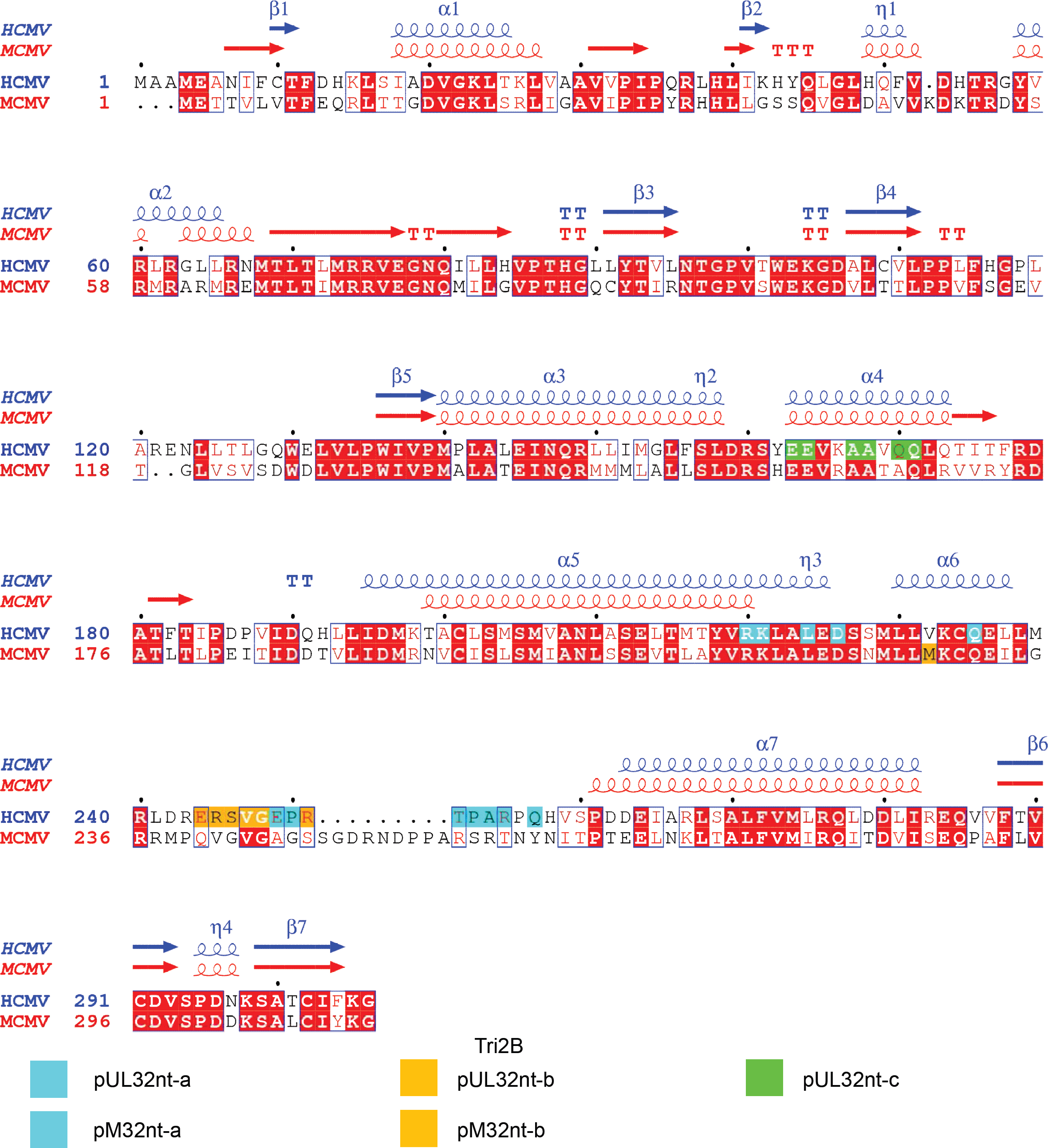
Secondary structure and sequence alignment for Tri2B in MCMV and HCMV. Schematic representations of the amino acid sequence and secondary structure alignment for Tri2B in HCMV and MCMV analyzed and displayed by *ESPript 3.0*. Arrow and spiral represent β-sheet and α-helix, respectively. Residues in Tri2B interacting with pUL32-a/pM32-a (cyan), pUL32-b/pM32-b (orange red), and pUL32-c (green, only in HCMV) are colored.

**FIG. S15.**
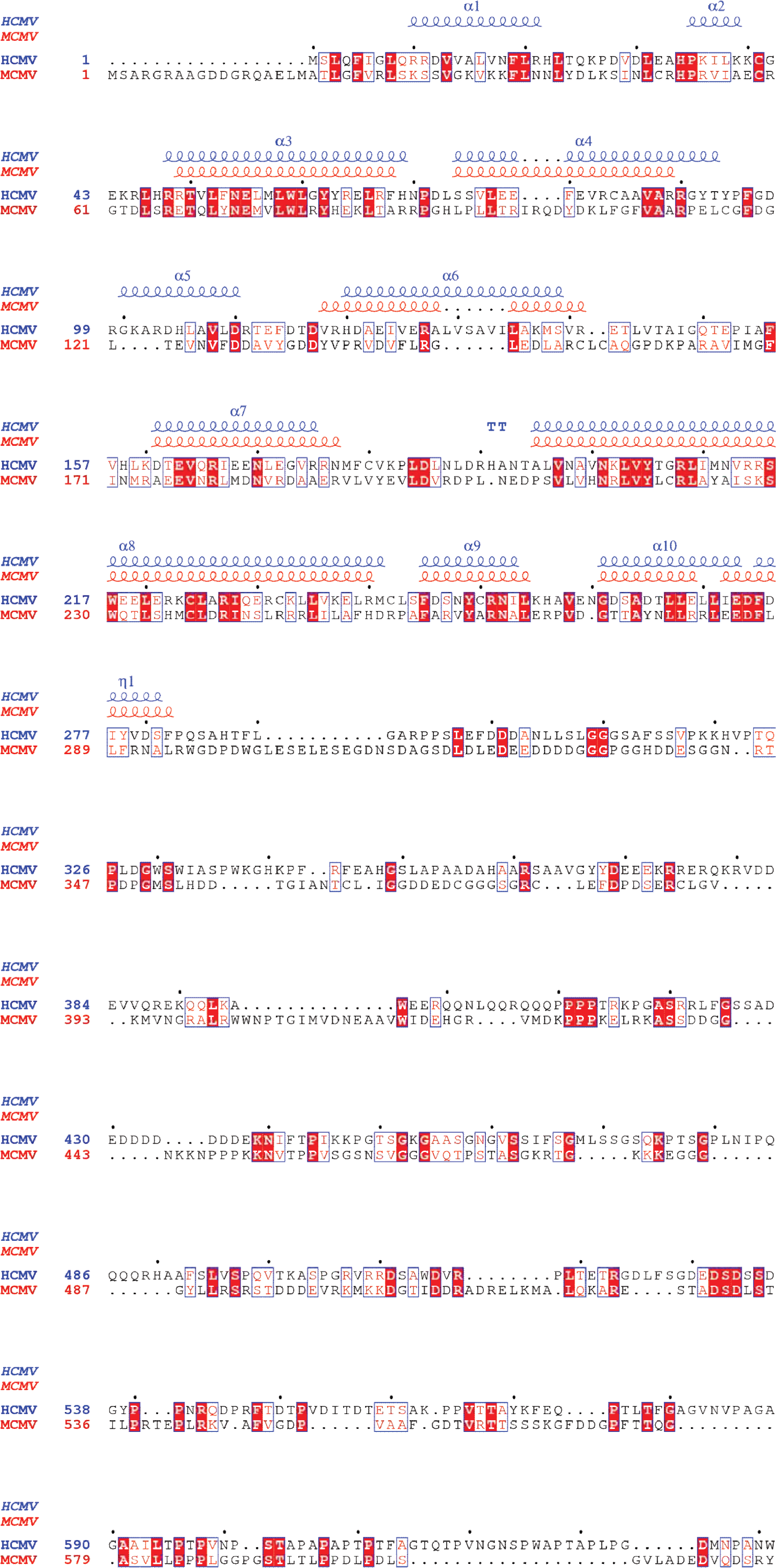
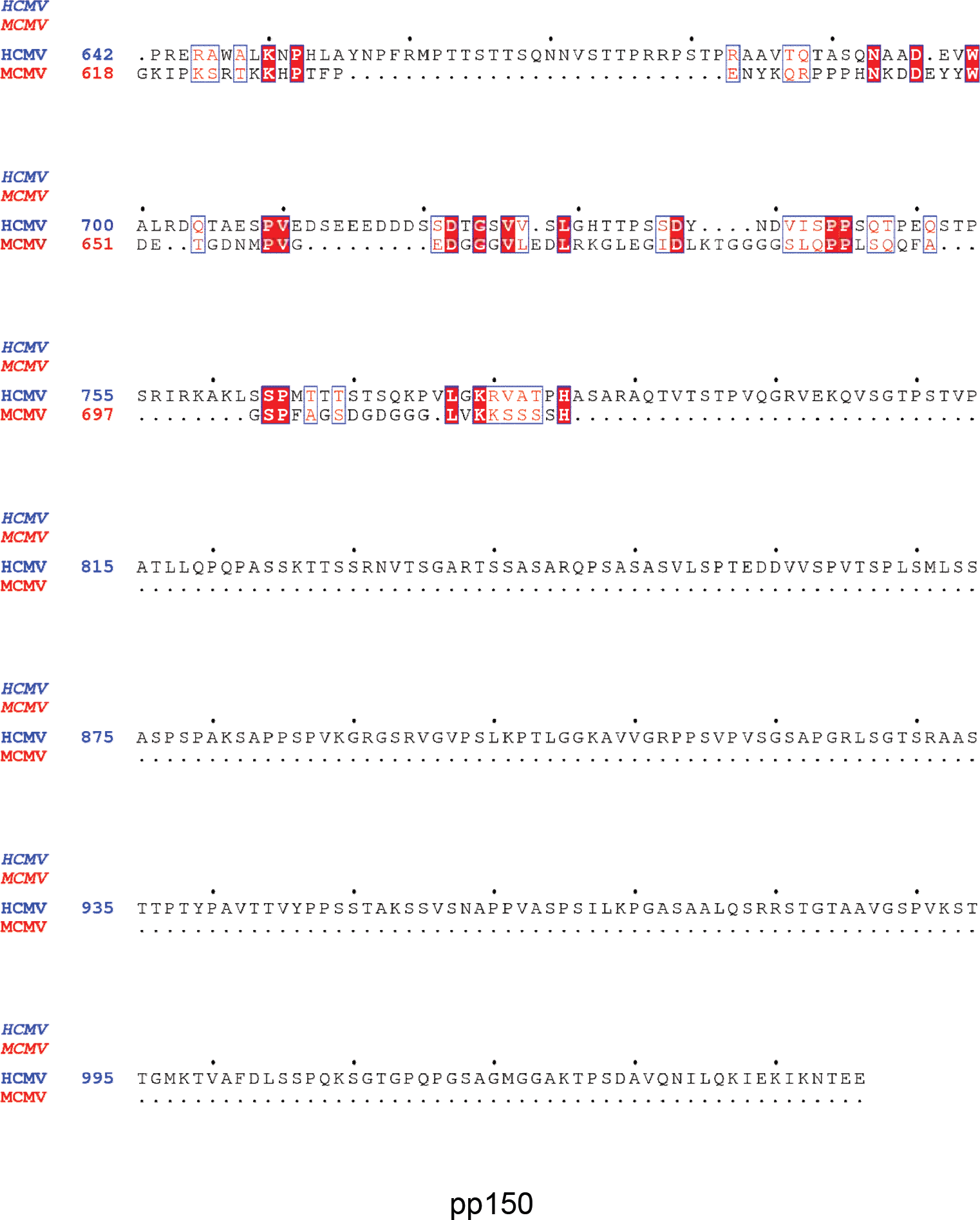
Secondary structure and sequence alignment for pM32 and pUL32 in MCMV and HCMV. Schematic representations of the amino acid sequence and secondary structure alignment for pUL32nt and pM32nt in HCMV and MCMV analyzed and displayed by *ESPript 3.0.* Spiral represents α-helix.

## Supplementary movie legends

**Movie S1. Shaded surface view of the icosahedral reconstruction of the MCMV virion at 5Å resolution.** Only the capsid and associated tegument protein pM32 (pp150) are visible.

**Movie S2. High-resolution structure features in the 3.8-Å resolution sub-particle reconstruction from the region around the 2-fold axis.**

**Movie S3. High-resolution structure features in the 3.6-Å resolution sub-particle reconstruction from the region around the 3-fold axis.**

**Movie S4. High-resolution structure features in the 3.8-Å resolution sub-particle reconstruction from the region around the 5-fold axis.**

## References

1. Gilbert C, Boivin G. 2005. Human Cytomegalovirus Resistance to Antiviral Drugs. Antimicrobial Agents and Chemotherapy 49:873–883.

2. Lemmermann NAW, Reddehase MJ. 2016. Refining human T-cell immunotherapy of cytomegalovirus disease: a mouse model with ‘humanized’ antigen presentation as a new preclinical study tool. Medical Microbiology and Immunology 205:549–561.

3. Pan D, Han T, Tang S, Xu W, Bao Q, Sun Y, Xuan B, Qian Z. 2018. Murine Cytomegalovirus Protein pM91 Interacts with pM79 and Is Critical for Viral Late Gene Expression. Journal of Virology 92.

4. Bootz A, Karbach A, Spindler J, Kropff B, Reuter N, Sticht H, Winkler TH, Britt WJ, Mach M. 2017. Protective capacity of neutralizing and non-neutralizing antibodies against glycoprotein B of cytomegalovirus. PLOS Pathogens 13:e1006601.

5. Mutnal MB, Hu S, Little MR, Lokensgard JR. 2011. Memory T cells persisting in the brain following MCMV infection induce long-term microglial activation via interferon-?. Journal of NeuroVirology 17:424.

6. Krmpotic A, Bubic I, Polic B, Lucin P, Jonjic S. 2003. Pathogenesis of murine cytomegalovirus infection. Microbes and Infection 5:1263–1277.

7. Loo CP, Snyder CM, Hill AB. 2017. Blocking Virus Replication during Acute Murine Cytomegalovirus Infection Paradoxically Prolongs Antigen Presentation and Increases the CD8<sup>+</sup> T Cell Response by Preventing Type I IFN–Dependent Depletion of Dendritic Cells. The Journal of Immunology 198:383.

8. Kern ER, Collins DJ, Wan WB, Beadle JR, Hostetler KY, Quenelle DC. 2004. Oral Treatment of Murine Cytomegalovirus Infections with Ether Lipid Esters of Cidofovir. Antimicrobial Agents and Chemotherapy 48:3516–3522.

9. Smee DF, Pease A, Burger RA, Huffman JH, Morrey JD, Okleberry KM, Noble RL, Sidwell RW. 1992. Ganciclovir Treatment of Murine Cytomegalovirus Infection in Mice Immunosuppressed by Prior Infection with Friend Leukaemia Virus. Antiviral Chemistry and Chemotherapy 3:327–333.

10. Hiršl L, Brizic I, Jenuš T, Juranic Lisnic V, Reichel JJ, Jurkovic S, Krmpotic A, Jonjic S. 2018. Murine CMV Expressing the High Affinity NKG2D Ligand MULT-1: A Model for the Development of Cytomegalovirus-Based Vaccines. Frontiers in Immunology 9.

11. Morello CS, Cranmer LD, Spector DH. 2000. Suppression of Murine Cytomegalovirus (MCMV) Replication with a DNA Vaccine Encoding MCMV M84 (a Homolog of Human Cytomegalovirus pp65). Journal of Virology 74:3696.

12. Dai X, Zhou ZH. 2018. Structure of the herpes simplex virus 1 capsid with associated tegument protein complexes. Science 360.

13. Dai X, Gong D, Lim H, Jih J, Wu T-T, Sun R, Zhou ZH. 2018. Structure and mutagenesis reveal essential capsid protein interactions for KSHV replication. Nature 553:521.

14. Yu X, Jih J, Jiang J, Zhou ZH. 2017. Atomic structure of the human cytomegalovirus capsid with its securing tegument layer of pp150. Science 356.

15. Close WL, Anderson AN, Pellett PE. 2018. Betaherpesvirus Virion Assembly and Egress. Advances in experimental medicine and biology 1045:167–207.

16. AuCoin DP, Smith GB, Meiering CD, Mocarski ES. 2006. Betaherpesvirus-conserved cytomegalovirus tegument protein ppUL32 (pp150) controls cytoplasmic events during virion maturation. J Virol 80:8199–8210.

17. Mocarski ES, Shenk T, Griffiths PD, Pass RF. 2013. Cytomegaloviruses, p. 1960–2014. *In* Knipe DM HP, Cohen JI et al (ed.), Fields Virology. Philadelphia, Pa.: Lippincott-William & Wilkins.

18. Baxter MK, Gibson W. 2001. Cytomegalovirus basic phosphoprotein (pUL32) binds to capsids in vitro through its amino one-third. J Virol 75:6865–6873.

19. Zhang X, Hong Zhou Z. 2011. Limiting factors in atomic resolution cryo electron microscopy: No simple tricks. Journal of Structural Biology 175:253–263.

20. Leong PA, Yu X, Zhou ZH, Jensen GJ. 2010. Chapter Fourteen - Correcting for the Ewald Sphere in High-Resolution Single-Particle Reconstructions, p. 369–380. *In* Jensen GJ (ed.), Methods in Enzymology, vol. 482. Academic Press.

21. Kucukelbir A, Sigworth FJ, Tagare HD. 2013. Quantifying the local resolution of cryo-EM density maps. Nature Methods 11:63.

22. Rochat RH, Liu X, Murata K, Nagayama K, Rixon FJ, Chiu W. 2011. Seeing the portal in herpes simplex virus type 1 B capsids. J Virol 85:1871-1874.

23. Deng B, O’Connor CM. Kedes DH. Zhou ZH. 2007. Direct visualization of the putative portal in the Kaposi’s sarcoma-associated herpesvirus capsid by cryoelectron tomography. J Virol 81:3640-3644.

24. Chang JT, Schmid MF, Rixon FJ, Chiu W. 2007. Electron cryotomography reveals the portal in the herpesvirus capsid. J Virol 81:2065-2068.

25. Cardone G, Winkler DC, Trus BL, Cheng N, Heuser JE, Newcomb WW, Brown JC, Steven AC. 2007. Visualization of the herpes simplex virus portal in situ by cryo-electron tomography. Virology 361:426-434.

26. Zhou ZH, Hui Wong H, Shah S, Jih J, O’Connor Christine M, Sherman Michael B. Kedes Dean H, Schein S. 2014. Four Levels of Hierarchical Organization, Including Noncovalent Chainmail, Brace the Mature Tumor Herpesvirus Capsid against Pressurization. Structure 22:1385-1398.

27. Chen DH, Jiang H, Lee M, Liu F, Zhou ZH. 1999. Three-dimensional visualization of tegument/capsid interactions in the intact human cytomegalovirus. Virology 260:10-16.

28. Messerle M. Crnkovic I, Hammerschmidt W, Ziegler H, Koszinowski UH. 1997. Cloning and mutagenesis of a herpesvirus genome as an infectious bacterial artificial chromosome. Proc Natl Acad Sci U S A 94:14759-14763.

29. Dunn W, Chou C, Li H, Hai R. Patterson D, Stoic V, Zhu H, Liu F. 2003. Functional profiling of a human cytomegalovirus genome. Proc Natl Acad Sci U S A 100:14223-14228.

30. Wagner M, Jonjic S, Koszinowski UH, Messerle M. 1999. Systematic excision of vector sequences from the BAC-cloned herpesvirus genome during virus reconstitution. J Virol 73:7056–7060.

31. Feng L, Sheng J, Vu G-P, Liu Y, Foo C, Wu S, Trang P, Paliza-Carre M, Ran Y, Yang X, Sun X, Deng Z, Zhou T, Lu S, Li H, Liu F. 2018. Human cytomegalovirus UL23 inhibits transcription of interferon-? stimulated genes and blocks antiviral interferon-? responses by interacting with human N-myc interactor protein. PLOS Pathogens 14:e1006867.

32. Zhou ZH, Dougherty M, Jakana J, He J, Rixon FJ, Chiu W. 2000. Seeing the herpesvirus capsid at 8.5 Å. Science 288:877–880.

33. Wu L, Lo P, Yu X, Stoops JK, Forghani B, Zhou ZH. 2000. Three-dimensional structure of the human herpesvirus 8 capsid. J Virol 74:9646–9654.

34. Trus BL, Gibson W, Cheng N, Steven AC. 1999. Capsid structure of simian cytomegalovirus from cryoelectron microscopy: evidence for tegument attachment sites. J Virol 73:2181–2192.

35. Zhou ZH, Stoops JK, Krueger GRF. 2006. Ultrastructure and Assembly of Human Herpesvirus-6 (HHV-6), p. 11–21. *In* Krueger G, Ablashi D (ed.), Perspectives in Medical Virology, vol. 12. Elsevier.

36. Dai X, Yu X, Gong H, Jiang X, Abenes G, Liu H, Shivakoti S, Britt WJ, Zhu H, Liu F, Zhou ZH. 2013. The Smallest Capsid Protein Mediates Binding of the Essential Tegument Protein pp150 to Stabilize DNA-Containing Capsids in Human Cytomegalovirus. PLOS Pathogens 9:e1003525.

37. Mindell JA, Grigorieff N. 2003. Accurate determination of local defocus and specimen tilt in electron microscopy. Journal of Structural Biology 142:334–347.

38. Kivioja T, Ravantti J, Verkhovsky A, Ukkonen E, Bamford D. 2000. Local Average Intensity-Based Method for Identifying Spherical Particles in Electron Micrographs. Journal of Structural Biology 131:126–134.

39. Ludtke SJ, Baldwin PR, Chiu W. 1999. EMAN: Semiautomated Software for High-Resolution Single-Particle Reconstructions. Journal of Structural Biology 128:82–97.

40. Liang Y, Ke EY, Zhou ZH. 2002. IMIRS: a high-resolution 3D reconstruction package integrated with a relational image database. J Struct Biol 137:292–304.

41. Liu H, Cheng L, Zeng S, Cai C, Zhou ZH, Yang Q. 2008. Symmetry-adapted spherical harmonics method for high-resolution 3D single-particle reconstructions. J. Struct. Biol. 161:64–73.

42. Zhang X, Zhang X, Hong Zhou Z. 2010. Low cost, high performance GPU computing solution for atomic resolution cryoEM single-particle reconstruction. Journal of Structural Biology 172:400–406.

43. Ilca SL, Kotecha A, Sun X, Poranen MM, Stuart DI, Huiskonen JT. 2015. Localized reconstruction of subunits from electron cryomicroscopy images of macromolecular complexes. Nature Communications 6:8843.

44. Scheres SHW. 2012. RELION: Implementation of a Bayesian approach to cryo-EM structure determination. Journal of Structural Biology 180:519–530.

45. Rosenthal PB, Henderson R. 2003. Optimal Determination of Particle Orientation, Absolute Hand, and Contrast Loss in Single-particle Electron Cryomicroscopy. Journal of Molecular Biology 333:721–745.

46. Pettersen EF, Goddard TD, Huang CC, Couch GS, Greenblatt DM, Meng EC, Ferrin TE. 2004. UCSF Chimera—A visualization system for exploratory research and analysis. Journal of Computational Chemistry 25:1605–1612.

47. Schwede T, Kopp Jr, Guex N, Peitsch MC. 2003. SWISS-MODEL: an automated protein homology-modeling server. Nucleic Acids Research 31:3381–3385.

48. Emsley P, Cowtan K. 2004. Coot: model-building tools for molecular graphics. Acta Crystallographica Section D 60:2126–2132.

49. Afonine PV, Grosse-Kunstleve RW, Echols N, Headd JJ, Moriarty NW, Mustyakimov M, Terwilliger TC, Urzhumtsev A, Zwart PH, Adams PD. 2012. Towards automated crystallographic structure refinement with phenix.refine. Acta crystallographica. Section D, Biological crystallography 68:352–367.

50. Kelley LA, Mezulis S, Yates CM, Wass MN, Sternberg MJE. 2015. The Phyre2 web portal for protein modeling, prediction and analysis. Nature Protocols 10:845.

51. Abenes G, Chan K, Lee M, Haghjoo E, Zhu J, Zhou T, Zhan X, Liu F. 2004. Murine cytomegalovirus with a transposon insertional mutation at open reading frame m155 is deficient in growth and virulence in mice. Journal of Virology 78:6891–6899.

52. Jiang X, Gong H, Chen Y-C, Vu G-P, Trang P, Zhang C-Y, Lu S, Liu F. 2012. Effective inhibition of cytomegalovirus infection by external guide sequences in mice. Proceedings of the National Academy of Sciences 109:13070.

53. Zhou ZH, Chen DH, Jakana J, Rixon FJ, Chiu W. 1999. Visualization of tegument-capsid interactions and DNA in intact herpes simplex virus type 1 virions. J Virol 73:3210–3218.

54. Suloway C, Pulokas J, Fellmann D, Cheng A, Guerra F, Quispe J, Stagg S, Potter CS, Carragher B. 2005. Automated molecular microscopy: The new Leginon system. J Struct Biol 151:41–60.

